# The Polycomb-dependent epigenome controls β-cell dysfunction, dedifferentiation and diabetes

**DOI:** 10.1101/205641

**Authors:** Tess Tsai-Hsiu Lu, Steffen Heyne, Erez Dror, Eduard Casas, Laura Leonhardt, Thorina Boenke, Chih-Hsiang Yang, Sagar, Laura Arrigoni, Kevin Dalgaard, Raffaele Teperino, Lennart Enders, Madhan Selvaraj, Marius Ruf, Sunil Jayaramaiah Raja, Huafeng Xie, Ulrike Boenisch, Stuart H. Orkin, Francis C Lynn, Brad G. Hoffman, Dominic Grün, Tanya Vavouri, Adelheid Lempradl, J. Andrew Pospisilik

## Abstract

Chromatin is the physical template that stabilizes and specifies transcriptional programs. To date, it remains largely unclear to what extent chromatin machinery contributes to the susceptibility and progression of complex diseases. Here, we combined deep epigenome mapping with single cell transcriptomics to mine for evidence of chromatin dysregulation in type-2 diabetes. We identify two chromatin-state signatures that track the trajectory of β-cell dysfunction in mice and humans: ectopic activation of bivalent Polycomb-domains and a loss of expression at a subclass of highly active domains containing key lineage-defining genes. β-cell specific deletion of Polycomb (Eed/PRC2) triggers parallel transcriptional signatures. Intriguingly, these β-cell Eed-knockouts also exhibit highly penetrant hyperglycemia-independent dedifferentiation indicating that Polycomb dysregulation sensitizes the β-cell for dedifferentiation. These findings provide novel resources for exploring transcriptional and epigenetic control of β-cell (dys)function. They identify PRC2 as necessary for long-term maintenance of β-cell identity. The data suggest a two-hit model for loss of β-cell identity in diabetes and highlight epigenetic therapeutic potential to block dedifferentiation.

## INTRODUCTION

Complex genetic diseases such as cancer, autoimmunity, obesity and diabetes represent some of the greatest socio-economic challenges of our day. They result from predisposition set by many genetic susceptibility loci. Also, strong non-genetic components, often termed ‘environmental influences’, alter susceptibility, reversibility and triggering of disease. These non-genetic factors are believed to be mediated in part via chromatin, the macromolecular complex that packages and organizes cis-regulatory elements and transcription in the nucleus. To date, how chromatin regulation contributes to the etiologies of most complex diseases remains poorly understood.

Diabetes affects more than 400 million individuals worldwide (World Health Organization (WHO), 2016). It presents in numerous forms, including early-onset autoimmune type-1 diabetes and a multifactorial type-2 diabetes (T2D) associated with obesity, inflammation and insulin resistance. Ultimately, diabetes results from a lack of sufficient insulin secreting β-cell mass, an etiology that can result from impaired function, increased cell death, or loss of cell identity. The last decades have seen major advances in our understanding of β-cell identity, development, and plasticity. These notions embody transcriptional programming, a process in which lineage-defining transcription factors and chromatin machinery modify the structure, accessibility, and positioning of thousands of genes and regulatory enhancers to coordinate expression of large numbers of loci (Graf and Enver, 2009) (Holmberg and Perlmann, 2012). β-cells are believed to be highly plastic (Stanger and Hebrok, 2013)(Ziv et al., 2013). Pioneering studies over the last decades have revealed networks of transcriptional and chromatin regulators that drive β-cell lineage development (Oliver-Krasinski and Stoffers, 2008) (Arda et al., 2013) (van Arensbergen et al., 2010)(van Arensbergen et al., 2013) (Bramswig et al., 2013) (Xu et al., 2011), and provide barriers against transdifferentiation or loss of β-cell identity(Talchai et al., 2012) (Dhawan et al., 2011) (Puri et al., 2013)(Taylor et al., 2013) (Gao et al., 2014) (Papizan et al., 2011) (Swisa et al., 2017) (Ediger et al., 2017), (Collombat et al., 2009; Gutiérrez et al., 2017).

To date, it remains poorly understood how transcriptional programs are stabilized over the long cellular lifespans that can be found in vivo. Since adult β-cells are both long-lived and highly plastic, mechanisms are thought to be in place to continuously reinforce and stabilize the terminally differentiated state (Cnop et al., 2011) (Szabat et al., 2012). Seminal studies in both wild-type (Guo et al., 2013) and in genetic (Talchai et al., 2012) (Brereton et al., 2014) (Wang et al., 2014) models have highlighted relative losses of β-cell identity, or ‘dedifferentiation’, with mounting metabolic stress. Dedifferentiation was coined to describe *either* a reversal of the differentiation trajectory back towards progenitor states *or* a loss of terminal differentiation markers and phenotypes (Holmberg and Perlmann, 2012)(Weir et al., 2013). Studies have documented the phenomenon in islet culture (Russ et al., 2008) and in T2D, in rodents and in humans tissues, and have focused on re-appearance of progenitor markers (ALDH1A; Cinti et al., 2016) and loss of lineage defining gene expression as cardinal features (PDX1, MAFA, NKX6-1, INS, and GLUT2; Guo et al., 2013). To date, aside from identification of a limited number of inducers (hyperglycemia, β-cell inexcitability, and FoxO1 deficiency), we have a little understanding of the molecular mechanisms that dictate how and when dedifferentiation occurs.

One chromatin regulatory system important to defining cell fate trajectories during development is Polycomb. The Polycomb system comprises two major sets of repressive complexes, PRC1 and PRC2, whose respective activities enable stable maintenance of gene silencing through time and cell-division (Bracken et al., 2006)(Margueron and Reinberg, 2011) (Schuettengruber and Cavalli, 2009). PRC1 and PRC2 are non-redundant, evidenced perhaps best by divergent loss-of-function phenotypes. PRC2 methylates the histone lysine residue H3K27 and is sufficient to silence gene expression (Ferrari et al., 2014) (Margueron and Reinberg, 2011). PRC1 ubiquitinates H2AK119 at PRC2 marked domains, promoting chromatin compaction and further silencing (Simon and Kingston, 2013). The existence of numerous PRC1 and PRC2 sub-complexes has emerged in the literature, revealing additional unexplored complexities, as well as redundancies within complexes, a prime example being the core PRC2 methyltransferases themselves, Ezh1 and Ezh2 (van Arensbergen et al., 2013) (Ezhkova et al., 2011) (Xie et al., 2014) (Schimmelmann et al., 2016).

Here, we used unbiased epigenome mapping and single-cell RNA-sequencing to explore the chromatin-dependence of transcriptional regulation in β-cells. We observed two progressive signatures of fundamental chromatin state-associated transcriptional dysregulation consistent between human type-2 diabetes and HFD-driven β-cell dysfunction, a loss-of-silencing at poised/bivalent Polycomb-domains, and, collapse of gene expression at a unique subset of highly-accessible active domains including many key lineage determinants. To test the relevance of Polycomb-dependent regulation, we generated β-cell-specific Eed-knockout mice to abrogate all PRC2 activities. Intriguingly, loss of Eed/PRC2 led to the activation of gene expression at bivalent domains and loss of transcription at a unique subset of highly-accessible active domains, a complete recapitulation of the transcriptional dysregulation observed in human and mouse β-cell dysfunction. Furthermore, loss of Eed ultimately triggered highly penetrant, hyperglycemia-independent dedifferentiation throughout the β-cell compartment. These findings demonstrate that Eed/PRC2 is required for global gene silencing and cell identity maintenance in terminally differentiated cells. They identify one of the first chromatin-based mechanisms required for stabilizing long-term β-cell identity and suggest core chromatin-regulation as one arm of a two-hit model of β-cell dedifferentiation.

## RESULTS

### Chromatin state specific dysregulation is a hallmark of β-cell dysfunction

In order to search for evidence of potential chromatin-driven regulatory events in β-cell dysfunction we generated and intersected two orthogonal genomic analyses (schematic; Figure 1A). First, we used ChIP-sequencing to map high-dimensional epigenomes of mouse pancreatic β-cells from young adult C57Bl6/J mice (8-12 weeks of age). We profiled histone marks characteristic for active and poised promoters (H3K4me3), enhancers (H3K27ac/H3K4me1), and transcribed coding regions (H3K36me3 and H3K27me1); heterochromatic- and Polycomb-silenced domains (H3K9me3; and H3K27me3/H2AK119Ub respectively); quiescent intergenic regions (H3K27me2); transcription and accessibility (RNA-pol2); and complemented these with measurements of DNA-methylation, an epigenetic mark which correlates depending on context with transcription, accessibility, CG-density and/or promoter-silencing (WGBS; Avrahami et al., 2015). This extensive dataset provides in-depth information on the nature of chromatin and transcriptional-state in β-cells genome-wide, including at cis-regulatory elements, highly active loci (Nkx6-1; Figure 1B), silent genes (e.g. Vmn1r115; Figure 1C), and loci poised for transcriptional activation or repression (Cck; Figure 1D). These data represent a reference-level epigenome as defined by the international human epigenome consortium and a powerful resource for the β-cell and (epi)genomic communities.

**1.**
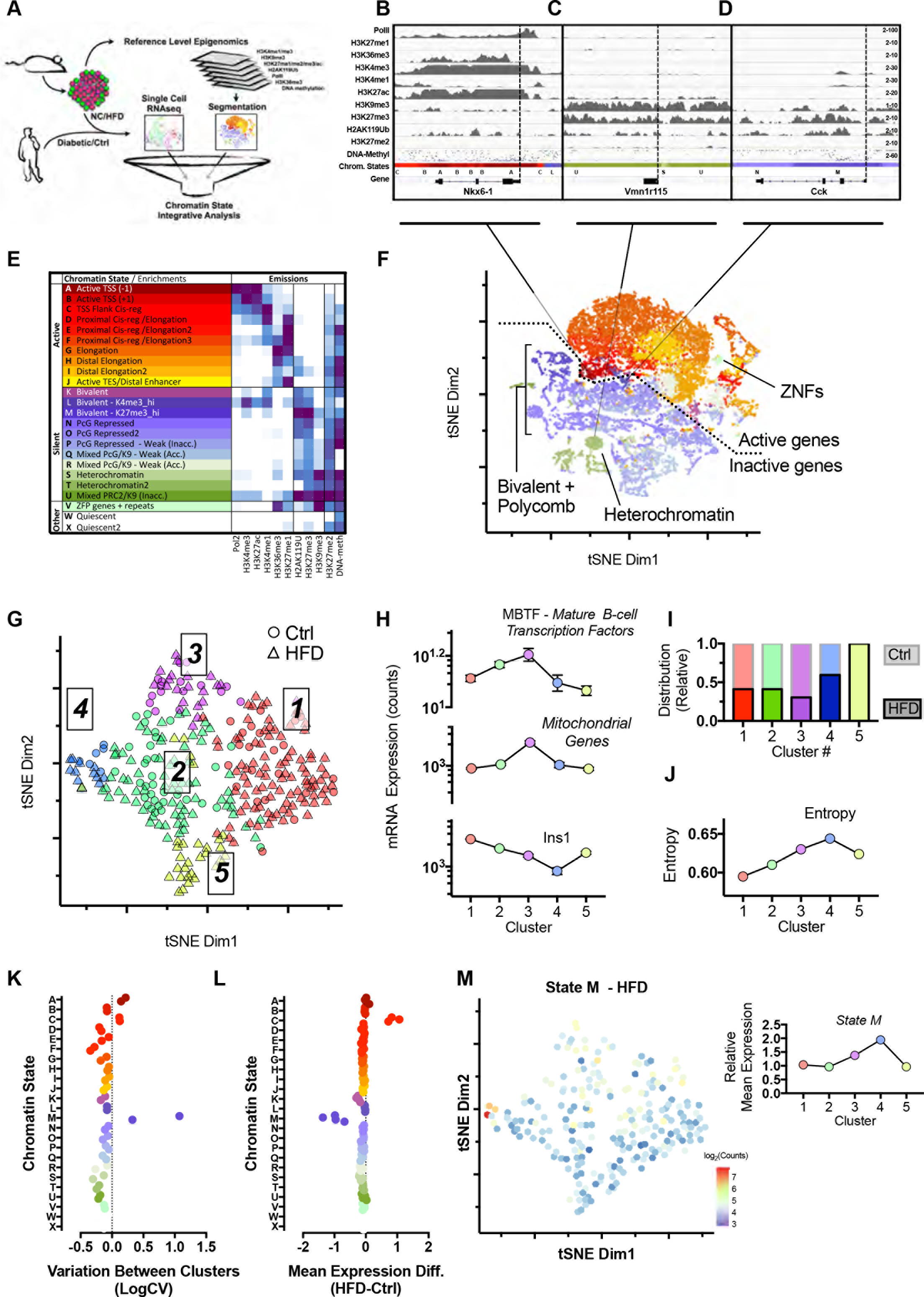
Chromatin state specific dysregulation is a hallmark of β Cell dysfunction. A. Schematic of the generation, intersection, and analyses of chromatin driven regulatory events in β Cell dysfunction. B-D. Genome browser view of histone mark occupancy and DNA-methylation at Nkx6-1 (State A, Fig 1B), Vmn1 r115 (State S, Fig. 1C) and Cck (State M, Fig 1D) gene loci. Dash line indicates TSS E. Histone mark characteristics in healthy islets. EpiCSeq chromatin state segmentation and manual annotation according to emissions and annotations in D. (Inacc=lnaccessible; +1 or −1=00B1 nucleosome position around TSS) F. tSNE representation of epigenomic classes of all genes highlighting chromatin states in annotated colours based on Figure. 1E. Dotted line segregates active and inactive genes. Solid lines point toward the epigenomic cluster of Nkx6-1, Vmn1r115 and Cck gene respectively. G. tSNE representation of control diet (CTRL) and high-fat diet (HFD) β-cell transcriptomes clustering, indicating 5 distinct β Cell sub-types. H. Average expression of mature β Cell transcription factors (MβTF, top), mitochondrial genes (middle), and insulin (Ins1, bottom) from clusters 1 to 5. I. Relative fraction of cells across the clusters from either control (Ctrl) or high-fat diet (HFD) treated samples J. Median entropy level (Grün et al., 2016) across clusters 1 to 5 K. Variation in mean chromatin state gene expression between the 4 (Ctrl) or 5 (HFD) β-cell clusters for all chromatin states. CV=coefficient of variation. L. Difference in mean chromatin state gene expression for each of Clusters 1 to 4 between HFD and Ctrl for all chromatin states. M. tSNE representation of mean state M gene expression levels in high-fat diet (HFD) β-cell transcriptomes (left panel). Relative mean expression of State M genes across clusters 1 to 5 (right panel)

To simplify the high-dimensional data for flexible use for instance with GWAS or RNA-seq, we performed chromatin-state segmentation. Analogous to clustering or PCA algorithms, chromatin state segmentation reduces high-dimensional epigenome information into single chromatin ‘states’ providing every bin of the genome a single classification that summarizes the many chromatin attributes of the locus (Mammana and Chung, 2015)(Figure 1E-F and 1B-D colored bar for example loci; Figure S1A and S1B). Our 25 state segmentation partitioned all ~25,000 genes into highly transcribed (Nkx6.1; Figure 1B), repressed (Vmn1r115; Figure 1C), and poised (Cck; Figure 1D) compartments and thus established a flexible and unbiased platform for exploring β-cell chromatin function from the single gene to transcriptome-wide scales. The analysis and visual representation provide an unprecedented view of the heterogeneity of gene expression architecture in the β-cell nucleus. A wide range of distinct epigenomic classes of active and silent genes is highlighted by the tSNE (Figure 1F). The chromatin state segregation faithfully identified euchromatic and heterochromatic domains (Figure 1E and 1F), various classes of active, silenced and poised gene states (Fig. 1E and 1F and Figures S1A and S1B), cis-regulatory elements such as enhancers and promoters (Figure S1A), as well as megabase scale spatial organization of the genome (Figure S1C). Noteworthy given high interest in DNA methylation, the analysis for the first time successfully integrates DNA methylation into such segmentation, extending the utility of these algorithms to incorporate covalent modifications of DNA. Thus, we generated an integrated β-cell epigenome useful for unbiased epigenome exploration by all β-cell biologists.

In parallel, we used single cell RNA-sequencing (scRNA-seq) to characterize the dynamics of β-cell transcriptome heterogeneity in health and disease. Using a 384-well based CELseq2 pipeline, we generated high-quality transcriptomes from individual β-cells derived from either chow (NCD) or age-matched high-fat diet treated (HFD, 18 weeks) C57Bl6/J mice (Figure 1 G-J). The methodology combines FACS sorting with parallelized barcoding reactions, and importantly, unique molecular identifiers (UMI) to eliminate PCR-duplication biases that can confound such data. After normalization and filtering the dataset included ~300 cells with a median ~15,000 unique transcript molecules per cell. K-medioids clustering revealed 5 major clusters of β-cell transcriptomes indicating at least 5 distinct β-cell sub-states or sub-types (Figure 1G). The majority of cells fell into two clusters characterized by robust expression of Mature β-cell Transcription Factors (MβTF’s)(Pdx1, Nkx6-1, Nkx2-2, Mafa, Pax6, Neurod1, Isl1, and Ucn3) and functional markers including insulin (Clusters 1 and 2; Figures 1G and 1H). Cells in Cluster 3 were consistent with recently described ‘hub cells’ exhibiting moderate insulin levels and high mitochondrial gene expression (Johnston et al., 2016). The two remaining clusters, 4 and 5, contained fewer cells, exhibited relatively low maturity and functional marker expression (Figure 1G and 1H). Cells from clusters 4 and 5 were primarily derived from HFD-challenged mice, indicating that HFD triggers a heterogeneous response in the islet, promoting a bias in the distribution of cells towards dysfunctional or dedifferentiated states (Figure 1G-I). Importantly, careful examination revealed that cells from healthy control islets also populated the poorly or de-differentiated cluster 4 (7%; 6 out of 78 control cells)(Figure 1I). This demonstrates that moderately dedifferentiated β-cell states exist even in healthy islets. Together, these data provide an unbiased snapshot of mouse β-cell heterogeneity and sub-type distribution in health and disease. They demonstrate that HFD triggers a shift in the β-cell population towards poorly differentiated cell states and provide a platform for examining the progression of β-cell transcriptome dysregulation in high-resolution.

In a third step, we intersected the two orthogonal datasets. We grouped all genes by their chromatin state and looked for evidence of abnormal variation across the five functional β-cell clusters (Figure 1K-M). Overall, detectable gene expression showed minimal variation across most chromatin states (Fig. 1K) indicating that the β-cell epigenome is largely refractory to accumulating dysfunction. Intriguingly though, genes embedded in three states (A, C and M) did show signatures of chromatin state-associated dysregulation. The most striking variation was observed for State M, differing significantly between HFD and Control cells within the same clusters (Figure 1L), as well as differing substantially between functional clusters in each of the Control and HFD datasets (Figure 1K). Chromatin State M loci were ‘Bivalent’, characterized by high accessibility (DNA-methylation low), transcriptional quiescence (Pol2 and RNA-seq negative) and, importantly, by double transcriptional start site marking with the active promoter mark H3K4me3 and Polycomb-deposited H3K27me3 (Figure 1D and 1E). Careful examination revealed that bivalent State M genes - typically thought to be poised for activation or repression - increased expression as differentiation factors fell and entropy increased (Figure 1M). Gain of low-level gene expression (State M activation) and loss of highly expressed genes such as insulin, suggested elevated transcriptional entropy, or uniformity. High entropy is a transcriptome characteristic that tracks with ‘stemness’ as stem cells lack exceptionally active, cell-type specific expression programs. Calculation revealed increased entropy in the low functionality clusters and indicated Cluster 4 as the most dedifferentiated by unbiased genome-wide measure (Figure 1J) (Grün et al., 2016). These data corroborate the reductions in maturity and functional β-cell genes (Figure 1H). Thus, high fat diet triggers chromatin-state defined transcriptome dysregulation and loss of cell identity in pancreatic β-cells.

### Polycomb (dys)regulation and dedifferentiation in human type-2 diabetes

To extend these findings towards the human setting we looked for signs of comparable dysregulation in islet RNA-seq data generated from 11 diabetic (HbA1c > 6.5) and 51 non-diabetic (HbA1c < 6.0) islet donors (Fadista et al., 2014). We ordered transcriptomes by HbA1c and examined orthologue expression according to our 25 chromatin-state model (Figure 2A). The analysis revealed marked similarities to the mouse results. Most importantly, the genes in the bivalent state M (as well as the related Polycomb states L, N, and O) were upregulated in type-2 diabetic patients (Figure 2A). This finding was consistent with a general derepression or loss-of silencing genome-wide (Figure 2B; 462 up- versus 47 down-regulated genes). The Polycomb-state derepression was characterized by enriched alternate lineage factor expression, including genes characteristic of earlier stages in β-cell development (Figure 2C), as well as reduced expression of MβTF’s (Figure 2D). Both findings were true on both the single gene levels and when examining entire pathways previously defined to characterize mature (MA-R) or immature (IM-DM) compartments (Figures 2C-2E, Table S1). Finally, and consistent with loss of average β-cell identity, transcriptional entropy increased with loss of glucose control (Figure 2F). To rule out effects of using species-specific epigenome data, we also intersected publically available H3K27me3 and H3K4me3 ChIP-seq data from sorted human β-cells (Bramswig et al., 2013) with the same human type-2 diabetes islet RNA-seq datasets (Fadista et al., 2014). Consistent with the findings above, type-2 diabetes up-regulated genes were preferentially H3K27me3-marked in healthy islets, an enrichment signature that was maintained whether comparing to equally expressed or all genes as control (Figure 2G and 2H; Figure S2A). Thus, results from multiple human and mouse datasets indicate inappropriate activation of Polycomb-silenced bivalent domains, loss of transcription at key mature β-cell transcription factors, and elevated transcriptional entropy are novel conserved features of β-cell dysfunction.

**2.**
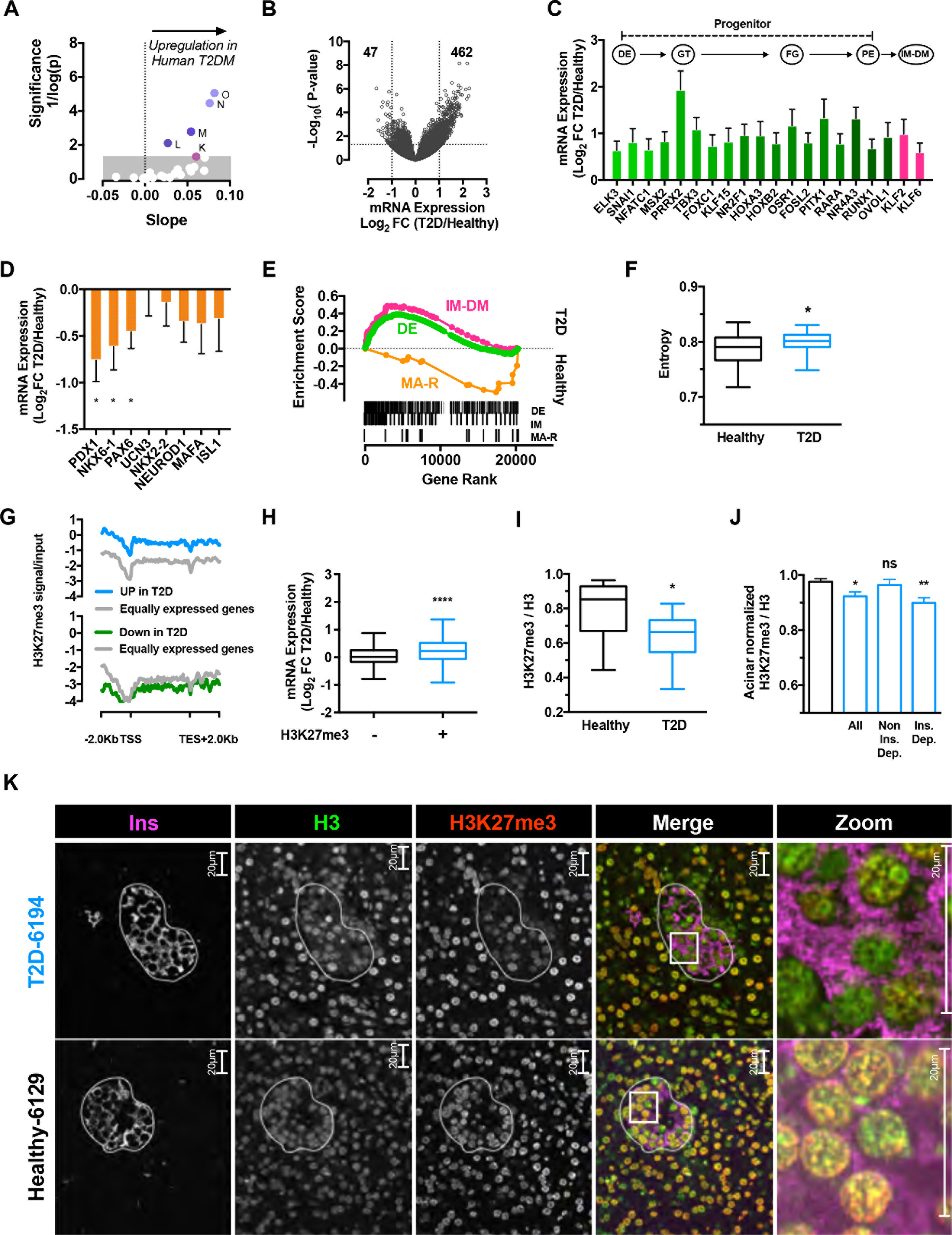
Polycomb (dys)regulation and dedifferentiation in human type-2 diabetes. A. Human orthologs embedded in bivalent polycomb repressed states (M) and related polycomb states (L, N and 0) are significantly and positively correlated with gene up-regulation in human T2D states (Fadista et al., 2014). Arrow indicates direction of positive correlation with gene up-regulation in human T2DM tested with linear regression. Upper boundary of grey box indicates p=0.05. B. A bias towards gene up-regulation in mRNA-seq data from human T2D islets. Volcano plot showing gene expression fold change between human T2D and healthy samples against p-value (−log10). Horizontal dotted line indicates p=0.05, vertical lines indicates ±2 fold difference. Number of significantly up or down regulated genes are highlighted. C. Mean expression fold change of transcription factors characteristic of developmental precursor stages (DE: definitive endoderm, GT: gut tube, FG: fore gut, PE: pancreatic endoderm; Xie et al., 2013) and immature β-cells (IM-DM; Blum et al., 2012). Arrow represent hypothetical developmental trajectory. All genes listed are core enrichment genes from GSEA analysis and significantly different between healthy and T2D islets with FDR-q<0.05. D. Mean expression fold change of Mature β-cell transcription factors between human T2D and healthy islet transcriptomes. * p<0.05 E. GSEA reveals enrichment of definitive endoderm (DE, Xie et al., 2013) and Immature (IM-DM, Blum et al., 2012) gene sets and depletion of mature β-cell genes (MA(R); REACTOME: *Regulation of gene expression in β-cell*) in human T2D islets FDR-q<0.05 for all gene sets shown. F. Increased median entropy in T2D islet vs healthy islet transcriptomes. Box plots whiskers indicate min to max. G. Average H3K27me3 signals across genes that are up or down-regulated in T2D, compared to genes that are equally expressed. TSS: transcription start site, TES: transcription end. H. Box-plot of H3K27me3 marked and non-marked gene expression changes reveals up-regulation of PRC2 target genes (H3K27me3+) in human T2D islets. I-J. Significant reduction of H3K27me3 immunostaining intensity in β-cells of pancreatic sections from T2D donors (n = 11, blue bars) relative to healthy donors (n = 11, black bars), normalizing to H3 intensity from the same nuclei (I) and when normalizing against same image acinar nuclei (J). T2D donors are combined (all) or stratified by insulin treatment dependency (Non ins. dep. or ins. dep.) K. Representative images of a pancreatic islet (outlined) from a healthy (left) and T2D (right) donor. Zoomed areas are outlined in boxes. Note the greener insulin positive nuclei in the T2D individual. All data represents mean ± SEM, unless otherwise stated. Box plots whisker indicates 1. 5 interquartile range, unless otherwise stated. * p < 0.05, ** p< 0.01, **** p<0.0001. ns= non-significant

To validate that the observations indeed resulted from a loss of Polycomb function, we compared H3K27me3 levels in human pancreas sections (nPOD) from 11 type-2 diabetic and 11 control BMI- and age-matched human islet donors. Sections were stained in unison, imaged in a double-blinded manner and quantitated using an unbiased CellProfiler pipeline. We quantified Insulin, the histone H3, and H3K27me3 in >30,000 cells covering at least 9 independent images per donor. Intriguingly, H3K27me3 staining was consistently reduced in diabetic β-cell nuclei (Figure 2I-K). Differences were maintained whether normalizing H3K27me3 to H3 as an internal reference (Figure 2I and 2K, reduced red/yellow nuclear staining within the islet boundary in T2D ‘Merge’), or to surrounding acinar cell nuclei to correct for staining heterogeneity (Figures 2J). Reductions in H3K27me3 were independent of BMI, age, C-peptide level and donor ethnicity (Figure S2B). Of note, insulin-dependent donors exhibited more substantial depletion of H3K27me3 than those on alternate therapies suggesting H3K27me3-association with disease severity (Figure 2J). Thus, human type-2 diabetes is characterized by a relative loss of PRC2 function.

### Eed/PRC2 deficiency triggers progressive β-cell dedifferentiation and diabetes

To genetically probe the consequences of loss of PRC2-silencing, we sought to generate β-cell specific PRC2 deficient mice. We first examined β-cell specific Ezh2 KO mice (βEzh2KO) previously reported to exhibit moderate diabetes secondary to defective β-cell proliferation (Chen et al., 2009). βEzh2KO animals were also mildly diabetic in our hands (not shown). Consistent with partial redundancy between Ezh1 and Ezh2 (Xie et al., 2014) and their co-expression in β-cells throughout development and into adulthood (not shown), βEzh2KO animals fail to completely lose PRC2 activity though they did exhibit transient, partial loss of H3K27me3 in the most rapidly dividing (5 weeks of age) growth phase (8 weeks of age)(Figure S3A and S3B). While this made the model unfit for our purpose, the findings highlight βEzh2KO mice as a highly unique example of the long-term consequences that can result from modest epigenetic dysregulation in early life. To abrogate all PRC2 activities we instead crossed the same Cre transgenic mice with mice bearing a deletable Eed allele (Figure 3A). Deletion of Eed disassembles both Ezh1- and Ezh2-containing complexes rendering both enzymes non-functional (Xie et al., 2014). β-cell-specific Eed knockout (βEedKO) mice were born healthy, at Mendelian ratios and indistinguishable from control (Ctrl) littermates. Efficient deletion was confirmed on the mRNA and DNA levels (Figure S3C and S3D). Complete loss of H3K27me2 and H3K27me3 immunoreactivity specifically in β-cells confirmed deletion on the functional level (Figure 3B). Thus, we generated β-cell specific Eed/PRC2 loss-of-function mice.

**3.**
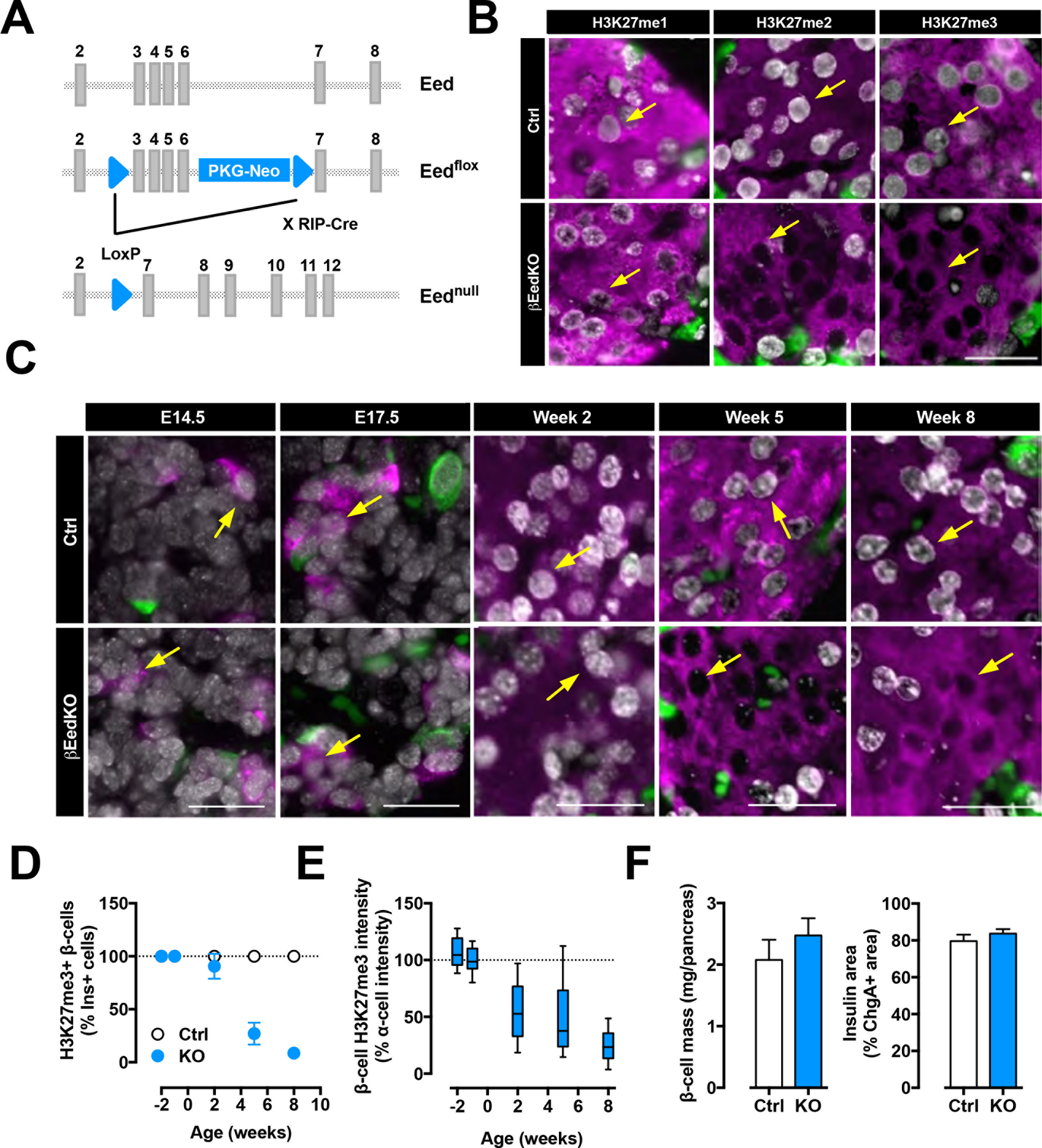
Eed/PRC2 deficiency triggers progressive β-cell dedifferentiation and diabetes. A. Schematic of the *Eed* targeting scheme. Light grey boxes depict exons (Xie et al., 2014). B. Immunofluorescence staining for H3K27me1, H3K27me2 and H3K27me3 (grey), insulin (magenta) and glucagon (green) in Ctrl and βEedKO showing robust loss of H3K27me2/3 specifically in β-cells. Yellow arrows indicate β-cell nuclei. C. Representative images for H3K27me3 staining (grey) in Ctrl and βEedKO showing progressive post-natal decline of the histone mark in β-cell with age. Insulin in magenta and glucagon in green. Yellow arrows indicate β-cell nuclei. D. Quantification of H3K27me3 positive β-cell number in βEedKO islets shows progressive and penetrant loss of H3K27me3. E. Mean β-cell H3K27me3 fluorescence signals in βEedKO islets shows progressive and penetrant loss of H3K27me3 signals with age. F. Quantification of total β-cell mass and insulin area per islet shows no significant difference between Ctrl and βEedKO animals. Data represents mean ± SEM of N=4-5 mice per genotype for each experiment. Scale bar = 25 uM.

Islet morphology in 8 week-old βEedKO animals was normal (Figure S3E). To determine the effective time-course of loss-of-function, we tracked H3K27me3 levels from late embryogenesis - when RIP-Cre becomes active (Xuan et al., 2002) - through to adulthood. H3K27me3 at E14.5 and E17.5 was unremarkable (Figure 3C, left two columns). After birth, however, β-cell H3K27me3 decreased progressively with a stark loss between 2 and 5 weeks of age (Figure 3C, middle and right columns). Loss of H3K27me3 was synchronized, affecting >90% of all insulin-positive cells by 8 weeks of age (Figure 3D and Figure 3E). Importantly, over the same age range we found no evidence of altered β-cell mass (Figure 3F), size or pancreatic insulin content (Figure S3F-G). Thus, β-cell Eed/PRC2 is dispensable for islet development and growth to adulthood.

To test the physiological consequences of β-cell specific PRC2 loss, we measured glucose tolerance in βEedKO animals at 8, 16 and 25 weeks of age. Eight-week old βEedKO mice were glucose tolerant (Figure 4A, top) with a tendency towards insulin over-secretion (Figures 4A, bottom). At 16 weeks of age βEedKO animals exhibited marked glucose intolerance (Figures 4B, top) and intriguingly, by 25 weeks, 100% of all knockouts exhibited overt diabetes (Figure 4C, top). Older knockouts exhibited strong and progressive reductions in both basal and glucose-stimulated insulin levels (Figure 4B-C, bottom) typically to undetectable levels. Without insulin-pellet supplementation all animals rapidly succumbed to the condition. Heterozygotes were unremarkable (Figure 4 A-C). The βEedKO phenotype was highly synchronized, with diabetes onset between 20-25 weeks of age in males (Figure S4A; 2hr fasting glucose >300mg/dl), and at ~40 weeks in females (not shown).

**4.**
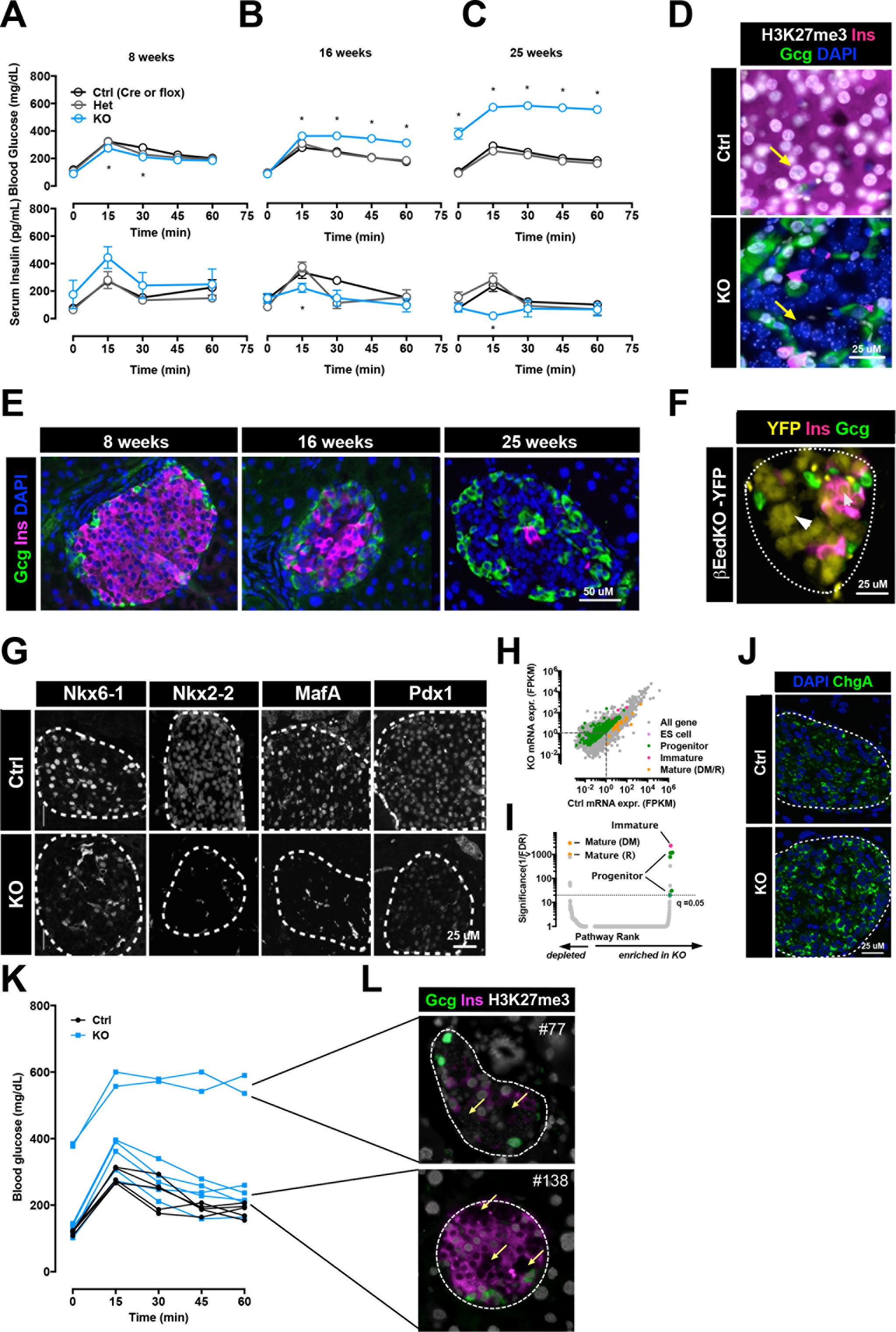
Eed/PRC2 deficiency triggers progressive β-cell dedifferentiation and diabetes. A. -C. βEedKO mice exhibit progressive glucose intolerance and diminished insulin secretory capacity during oral glucose tolerance test (1 g/kg) from 8 to 25 weeks of age. n=6-10 animals for each genotype. D. Hormone negative βEedKO islet cells are exclusively H3K27me3 negative. Yellow arrows indicate Ctrl and βEedKO cells harbouring or lacking H3K27me3 E. βEedKO islets exhibit progressive decline in insulin staining from 8 to 25 weeks of age F. Representative image of βEedKO islets harbouring a RIP-cre inducible YFP lineage tracer (yellow). Small white arrow indicates insulin-positive β-cells exhibiting YFP signals, and large white arrow indicates non-insulin expressing cells that also exhibits YFP signal. White dotted line outlines the islet for clarity. G. βEedKO islets exhibit loss of key β-cell markers (Pdx1, Mafa, Nkx2-2 and Nkx6-1). White dotted line outlines the islets. H. GSEA leading edge genes from “Progenitor” and “Immature” sets are up-regulated in βEedKO islets, whereas those from “Mature” gene sets are down-regulated (FDR<0.05). ES cell markers are not expressed. Dotted lines indicate FPKM=1 I. GSEA analysis of mRNA-seq expression data reveals significant enrichment of all “Progenitor” and “Immature” gene sets and depletion of “Mature” gene sets in βEedKO versus Ctrl islets. Dotted line indicates FDR-q<0.05 J. βEedKO islets exhibit robust immunoreactivity for Chromogranin-A K. Oral glucose tolerance test (1 g/kg) of 12 week old Pdx1-Cre;EedKO (n=6) or littermate control (n=5). Each curve represent a single mouse L. Representative image of the mosaic Eed/PRC2 loss-of-function deletion pattern in islets from the normoglycemic (#138, lower panel) and hyperglycemic (#77, upper panel) Pdx1-Cre;EedKO mice. Data represents mean ± SEM. * p<0.05-0.0001

Islets from 25-week old βEedKO mice were largely normal in size and morphology (Figure S4B). Organization and relative α−, δ− and PP-cell numbers were unremarkable. Intriguingly, the expected β-cell majority of the islet was devoid of hormones, staining negative for all major islet endocrine hormones including insulin (Figure 4D, 4E and Figure S4C). Consistent with onset of diabetes, βEedKO-driven loss of insulin immunoreactivity was synchronized and progressive between 8 and 25 weeks of age (Figure 4E). Hormone-negative βEedKO cells were all H3K27me3-negative, indicating they originated from the Eed-knockout β-cell compartment (Figure 4D). Lineage-tracing validated this notion. Using a RIP-Cre-inducible eYFP fluorescent reporter, the majority of hormone-negative βEedKO cells were eYFP reporter positive (Figure 4F, large arrow and Figure S4D). Thus, Eed/PRC2 is required for long-term maintenance of β-cell function, insulin expression, and glucose tolerance in vivo.

βEedKO cells exhibited reduced levels of the mature β-cell markers Pdx1, MafA, Nkx2-2 and Nkx6-1 (Figure 4G). Together with the complete loss of insulin expression, this finding indicated loss of β-cell identity. Interestingly, RNA-sequencing revealed very high correlation between wild type and βEedKO transcriptomes (R^2^>0.95; 25-weeks), indicating that despite the profound phenotype, knockout cells remained highly similar to healthy wild-type β-cells (Figure 4H). Deeper analysis revealed that βEedKO islets down-regulated mature β-cell genes as defined by the Melton group (Mature-DM) and Reactome (Mature-R; *Regulation of gene expression in /β-cell*) (Figure 4H and 4I; Table S2). At the same time, they up-regulated immature β-cell and progenitor specific genes (Figure 4H and 4I; Figure S4E; (Blum et al., 2012; Xie et al., 2013) indicating that cells had only undergone reversal of very late, terminal differentiation or maturation. Noteworthy, βEedKO islets increased expression of Gli2, a genetically-validated driver of dedifferentiation (Figure S4F; Landsman et al., 2011). They showed moderate reduction in FoxO1 and no marked regulation of Vhlh and Kcnj11 - genes previously shown to buffer loss of mature identity. No changes were observed in the earlier progenitor markers Sox17, Oct4, Sox9 or Ngn3 (Puri et al., 2013) (Talchai et al., 2012) (Wang et al., 2014) (Figures 4H, and Figure S4F). No evidence was found of altered cell death (TUNEL; Figure S4G) or marked β-to-α-cell transdifferentiation (by lineage tracing; not shown). Consistent with other diabetic models, we observed an ~15% increase in non-β-cell mass (Figure S4H). Importantly, hormone-negative βEedKO cells retained expression of Chromogranin A (Figure 4J), an indicator of maintained late endocrine character. Thus, loss of Polycomb function in β-cells results in specific loss of terminal identity.

Seminal studies highlighted hyperglycemia as one mechanistic driver of β-cell dedifferentiation. To address the contribution of hyperglycemia towards Eed/PRC2-dependent buffering of terminal identity, we used subcutaneous insulin implants to normalize blood glucose long-term in βEedKO mice, initiating treatment at moderate (~18 wks-of-age; normal fasting glucose) or late-stage pathology (~25 wks; >90% dedifferentiation). Interestingly, these interventions failed to abrogate the EedKO-triggered dedifferentiation (Figure S4i) suggesting that hyperglycemia is not an absolute requirement for Polycomb-associated dedifferentiation. In a separate experiment, we took advantage of the low deletion efficiency of the Pdx1-Cre transgenic line allowing the generation mice with a mosaic Eed/PRC2 loss-of-function deletion pattern in their islets. Pdx1-Cre;EedKO mice presented with a range of glycemic control from normoglycemic to moderate diabetes already at 12 weeks of age (Figure 4K). Importantly, all knockout mice, including normoglycemic individuals, exhibited cells showing loss of terminal differentiation. In all cases these cells were also Eed-deficient as evidenced by lack of H3K27me3 staining (Figure 4L). Together, these data indicate that dedifferentiation triggered by impaired PRC2 function is cell autonomous and does not absolutely require dysglycemia. They suggest that two independent (likely synergistic) axes contribute to dedifferentiation, one chromatin-associated, and the other coupled to hyperglycemia. Thus, Eed/PRC2 is required for long term maintenance of β-cell identity in vivo.

### Eed/PRC2 maintains global silencing in terminally differentiated β-cells

Exploring the transcriptional dysregulation more deeply, we observed changes recapitulating key aspects of our unbiased analysis of human and mouse β-cell dysfunction (Figure 5A and 5B). We observed ~5-fold more up- than down-regulation in βEedKO islets (Figure 5A). These changes were progressive and evident already at 8 weeks of age, prior to hyperglycemia and identity loss. This suggested a common underlying mechanism (Figure 5A, Table S3). Whereas down-regulation was observed primarily at active state A embedded genes (Figure 5B), up-regulation was biased towards the bivalent polycomb-silenced chromatin states L and M (DNA-methylation low, transcriptionally quiescent and H3K4me3/H3K27me3 double-positive)(Figure 5B). H3K27ac levels increased progressively after Eed knockout (Figure S5B). H3K27ac ChIP-seq revealed relatively dramatic ectopic H3K27ac deposition focussed at >10,000 sharp novel peaks (Figure 5C). These ectopic peaks were not randomly distributed, but rather appeared very specifically at pre-existing inactive but accessible promoters and enhancers as indicated by their precise intersection with H3K4me1 and H3K4me3 marked, DNA-methylation low loci (Figure 5D, areas shaded grey). Consistent with in vitro work by Ferrari et al. these data support the notion that histone acetylation and methylation activities compete for H3K27 residues at chromatin (Ferrari et al., 2014). It indicates that a similar competitive balance exists in vivo in terminally differentiated β-cells. Thus, Eed/PRC2 function is required to prevent inappropriate activation of bivalent, accessible genomic regions in terminally differentiated β-cells. The data highlight conditional β-cell Eed/PRC2 deletion as a model for the transcriptional dysregulation observed in human and mouse diabetes.

**5.**
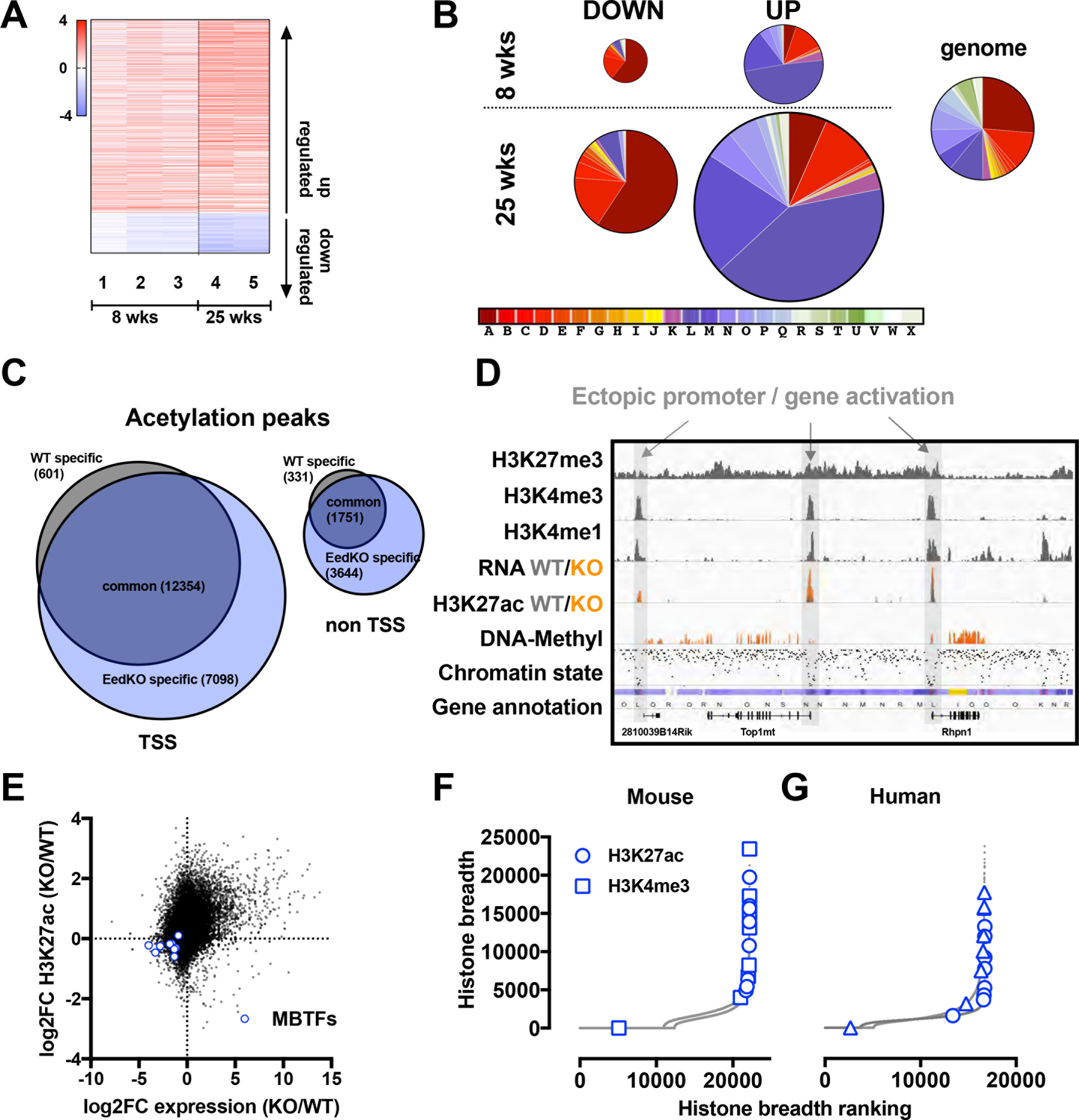
Eed/PRC2 maintains global transcriptional silencing in terminally differentiated β-cells. A. Clustered heatmap indicating 5 times more up- than down- regulation of βEedKO islets gene expression at both 8 and 25 weeks of age. Samples 1-5 refers to individual βEedKO RNA-seq samples (n=1-4 per sample) B. Pie chart showing chromatin state distribution of differentially expressed genes indicates an enrichment of States L and M (bivalent, polycomb-silenced) for up-regulated and State A for down-regulated genes at both 8 and 25 weeks of age. Far right panel shows chromatin state distribution of all genes. C. Venn diagram indicating gain of ectopic H3K27ac peaks in βEedKO genome at either TSS and non-TSS associated genomic loci. D. Genome browser view of ectopic H3K27ac peaks at accessible promoter and enhancer regions highlighted in blue. WT and KO tracks are overlaid for RNA and H3K27ac to facilitate for comparison. E. MβTFs exhibit specific depletion of H3K27ac marking and down-regulation of gene expression in βEedKO islets. F. -G. MβTFs exhibit broad H3K27ac and H3K4me3 signals and are amongst the top ranked genes according to H3K27ac and H3K4me3 breadth rankings in both mouse and human

### Eed/PRC2 protects transcription at epigenomically unique lineage genes

Notably, we observed a select few loci that were *depleted* of H3K27ac marking in KO islets despite the robust genome-wide deposition (Figure 5E). Consistent with the observed dedifferentiation, these loci included virtually all MβTF’s (Pdx1, Nkx6-1, Nkx2-2, Mafa, Pax6, Neurod1, Isl1, and Ucn3; Figure 5E). Interestingly, in our chromatin state analyses, these genes clustered extremely closely within the tSNE projection (Figure S5C). These data indicate for the first time to our knowledge that epigenome architecture at these β-cell-defining lineage genes is unique relative to the vast majority of active genes. Specifically, MβTF loci exhibited intense and broad active-mark deposition across the entirety of their loci (Figure 1B Nkx6-1; Figure 5F-G and S5A); they were devoid of silent marks and the genes themselves tended to be short, with few exons and little elongation mark accumulation (H3K36me3 / H3K27me1). Notably, they showed exceptionally broad and uniform Pol2 binding (Figure 1B) indicating distinct transcriptional control. Thus, Eed/PRC2 *protects* transcription at epigenomically unique lineage-defining loci.

### Ectopic transcription factor expression drives loss of cell identity in Min6 cells

A key signature found in both βEedKO and human type-2 diabetic islets was the ectopic expression of bivalent chromatin embedded transcription factors normally silenced by PRC2 in healthy islets. To test whether the ectopic activation of these genes was sufficient to drive the loss of cell identity, we identified 6 factors that were i. H3K27me3-marked (PRC2-regulated) in normal mouse islets, ii. H3K27me3-marked (PRC2-regulated) in normal human islets, iii. markedly up-regulated in both βEedKO and human type-2 diabetic islets. We co-transfected these 6 diabetes-specific, Polycomb-regulated *ectopic transcription factors* (ETFs)(Barx1, Hoxb7, Gata2, Pitx1, Twist1, and Zic1) into the β-cell line Min6-B1. After transfection we tracked gene expression responses in these non-cycling cells by single-cell RNA-sequencing, using vectorless and GFP-transfected cells as controls (schematic Figure 6A). We compared 359 single-cell transcriptomes including 50 control, 78 GFP and 231 ectopic transcription factor (ETF) over-expressing cells (Figure 6B). Unbiased clustering analysis revealed 4 clusters. Two were composed mainly of control and GFP transfected cells showed high insulin and MβTF transcription (Figure 6B and 6C, clusters 1 and 2). The remaining two were comprised of transfected cells exhibiting ETF over-expression and, importantly, loss of MβTF and *Ins1* mRNA expression, and thus, loss of β-cell identity (Figure 6B and 6C, Clusters 3 and 4). To understand this ETF-driven dedifferentiation, we ordered cells along the dedifferentiation trajectory using pseudotemporal ordering (Cluster 1, 2, 3, 4 in order). The unbiased ordering revealed two apparent phases of loss of identity. First, a very tight coupling was observed between increasing ETF expression and decreasing insulin expression. This was observed from ETF expression levels as low as 2-fold normal (Figure 6C; region between hashed lines). In a second phase, more substantial ETF expression tracked with a progressive loss of lineage defining MβTFs (Figure 6C, right of second hashed line). Also, ectopic TF overexpression also triggered down-regulation of the epigenomically unique class of H3K27ac-broad genes described above to include MβTFs (Figure 6C, bottom panel). Importantly, the levels of ectopic ETF expression found able to initiate insulin loss within this 3-day in vitro experiment were comparable to those observed in human diabetic islets observable even in bulk RNA-sequencing which merges expression profiles of functional and dysfunctional β-cells (Figure 6D). Interesting to note, hyperactivation of H3K27ac-broad genes was observed during the first phase. These data suggest existence of multiple important transcriptional networks that buffer cellular identity. This model provides a novel system for investigation such transcriptional dysregulation relevant to diabetes.

**6.**
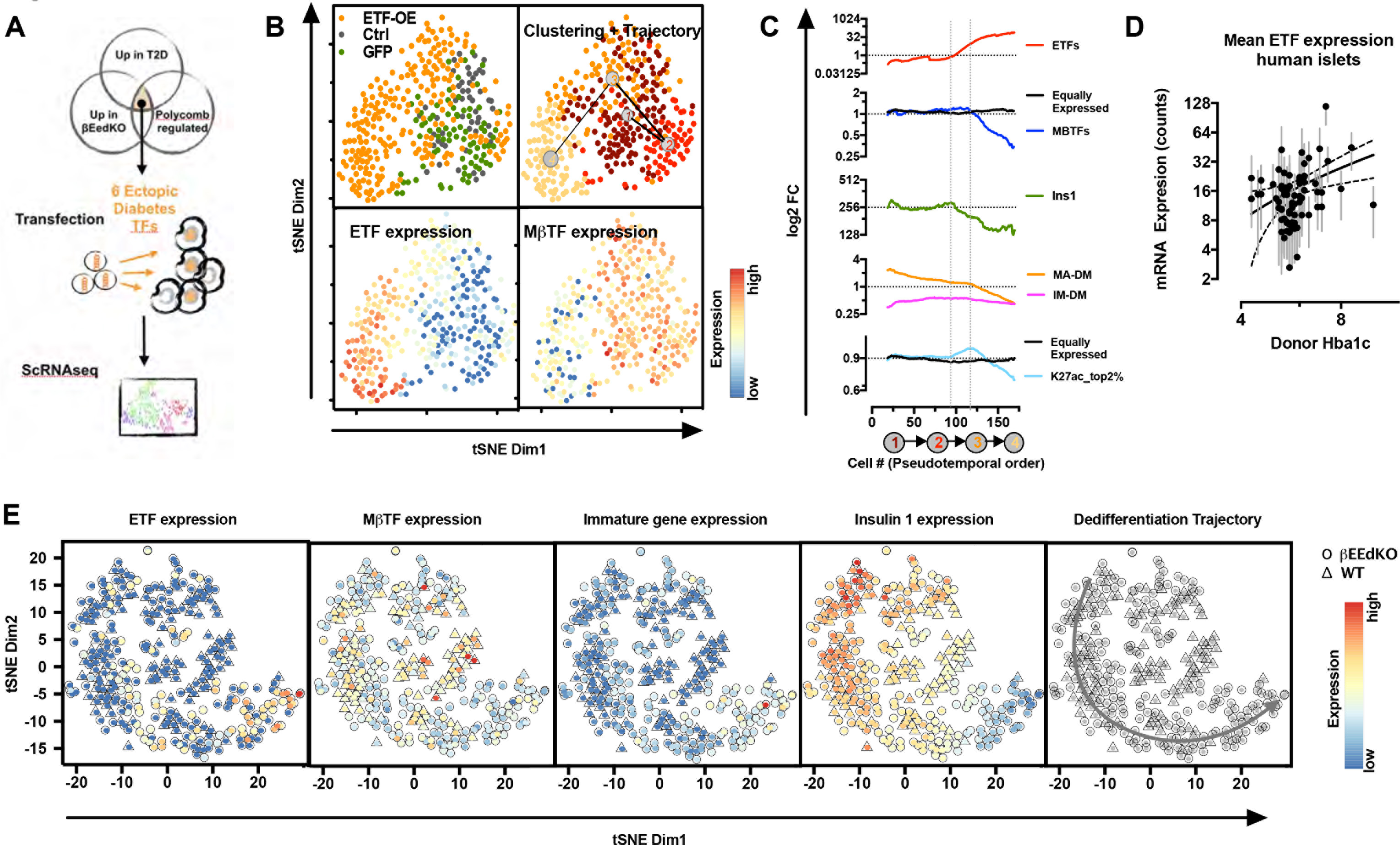
Ectopic transcription factor expression drives loss of cell identity in Min6 cells. A. Ectopic transcription factor expression-experiment design. B. t-SNE maps of control, GFP and ETF over-expressing cells from single-cell RNA sequencing experiment. Four clusters identified by RacelD2 are depicted with a red color gradient (Upper right panel). TF over-expressing dedifferentiated cells are enriched in cluster 4 and exhibit loss of MβTF expression C. Pseudo-temporal expression profiles along the dedifferentiation trajectory reveal down-regulation of MβTF, ‘Mature” and H3K27ac-broad genes upon ectopic TF over-expression. MA-DM: Mature gene set from Melton group, IM-DM: Immature gene set from Melton group (Blum et al., 2012). Note:H3K27ac top2% gene set excludes expression outliers Ins1 and Ins2. D. Mean expression of ETFs in human islets shows positive correlation with donor HbalC index. Linear regression (solid line) and 95% confidence interval (dash lines) are shown. E. t-SNE representation of the indicated gene(sets) in 14 week old βEedKO and WT cells. Data are from single-cell RNA sequencing experiment.

To further validate the in vivo relevance of these findings we searched for ETF overexpression in single cell RNA-sequencing data generated from 14 week old βEedKO β-cells (early in the progression towards dedifferentiation and diabetes). Ectopic ETF expression was only observed in knockout cells (Figure 6E), and even moderate expression was accompanied by loss of insulin and MβTF expression in those cells. All together, these findings argue for a model where ectopic bivalent transcription factor expression drives loss-of-cell-identity. Thus, ectopic activation of diabetes-specific ectopic transcripts is sufficient to drive β-cell dedifferentiation *in vitro*.

### PRC2-associated epigenome regulation is therapeutically targetable

Finally, we asked to what extent the observed Polycomb-dependent pathology might be amenable to intervention. To explore this we examined an *in vivo* scenario where we tested pharmacological administration of the histone deacetylase inhibitor, SAHA, in both and βEedKO-driven (Figure 7A) and HFD-driven (Figure 7B) pathologies. If etiology relies upon ectopic acetylation and transcriptional derepression at sensitive loci, we hypothesized that such an intervention should accelerate the dysfunction. Consistent with involvement of hyperacetylation and ectopic gene activation, SAHA treatment accelerated both HFD-induced and βEedKO-induced disease (Figure 7A and 7B) exacerbating glucose intolerance compared to non-SAHA treated controls., These data are consistent with the notion that ectopic acetylation and gene activation contribute towards β-cell dysfunction in vivo. They demonstrate that HFD- and EedKO-driven disease are sensitive to epigenetic interventions.

**7.**
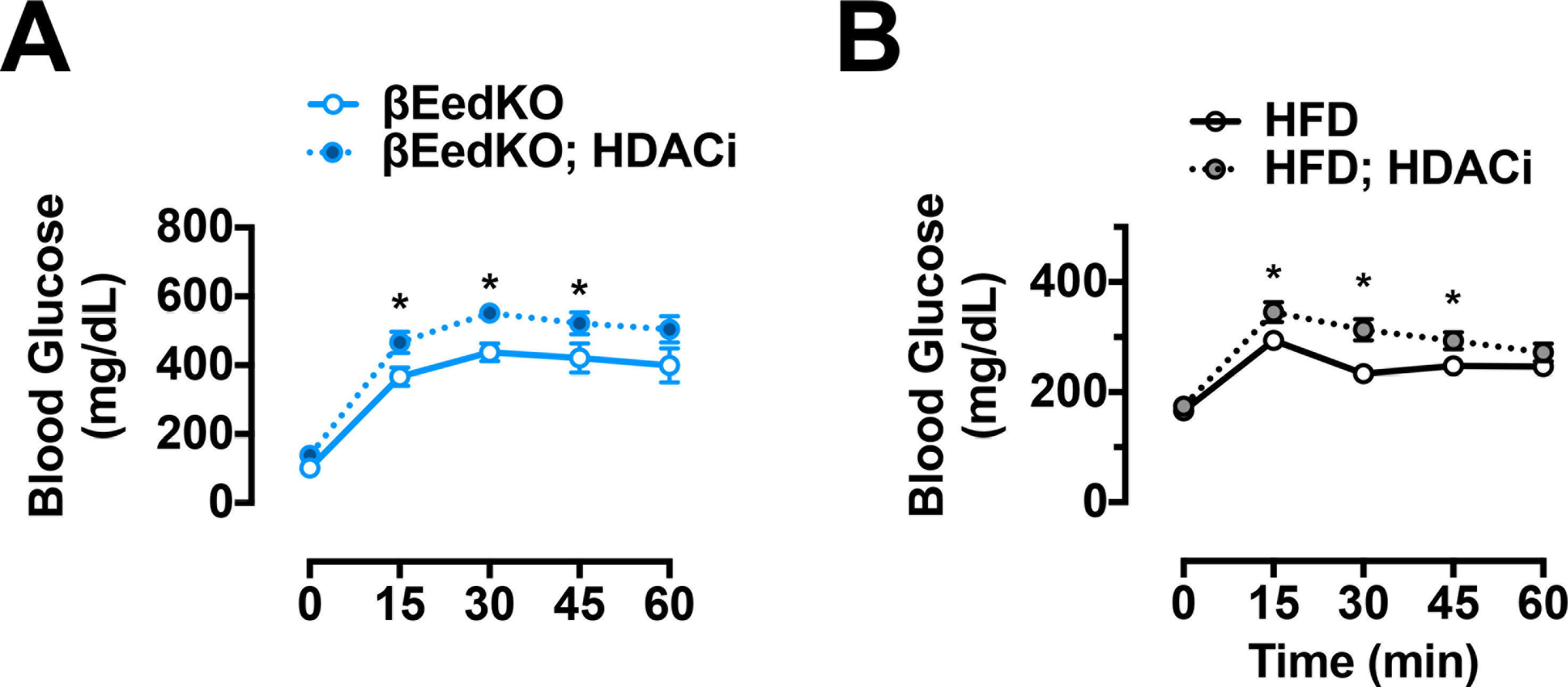
PRC2-associated epigenome regulation is therapeutically targetable. A. Oral glucose tolerance test (1 g/kg) of 25 week old βEedKO mice treated orally for 8 weeks with SAHA (2mg/day) or vehicle control (N=4-6 mice per group). B. Oral glucose tolerance test (1 g/kg) of 24 week old HFD-mice treated orally for 8 weeks with SAHA (2mg/day)or vehicle control (N=10 mice per group). Data are mean ± SEM. * p< 0.05

## DISCUSSION

Here, through the intersection of epigenome mapping and single-cell transcriptome analyses we identify chromatin state-specific loss of gene expression control as a novel feature of mouse and human beta-cell dysfunction. This dysregulation is associated with signatures of dedifferentiation. We provide first evidence that *Eed*/PRC2 is required for maintenance of global transcriptional silencing in differentiated pancreatic β-cells and that impairment of this regulatory axis is sufficient to drive progressive β-cell failure and dedifferentiation. These data, together with experiments demonstrating pharmacological (SAHA) manipulation of the process, as well as existence of comparable cells in normal healthy individuals, support a ‘two-hit’ model of beta-cell dedifferentiation, in which chromatin-dependent transcriptional dysregulation is a novel causal arm.

Ten years ago, Yamanaka and colleagues generated induced pluripotent stem cells by over-expressing a handful of transcription factors (Takahashi and Yamanaka, 2006). The experiment highlighted the complete plasticity potential locked within most differentiated cells and highlighted the robustness and specificity of mechanisms that maintain terminal cell identity. Indeed, the closer cells are to terminal differentiation, the more reinforced such mechanisms appear to be (Pasque et al., 2011). The analysis here, and in particular the tSNE visualization, highlight somewhat unappreciated epigenomic diversity at both active and silent genes, and in particular unique epigenome architecture at virtually all MβTF lineage factors. These highly active and broad regions likely relate to varying extents with previously described exceptionally broad H3K4me3 marked regions (Benayoun et al., 2014). Interestingly, our data identify these loci as amongst the most sensitive to transcriptional collapse during human and mouse β-cell decline and dedifferentiation. Indeed, loss of transcription at these loci must certainly contribute to dedifferentiation.

A number of studies have highlighted the importance of epigenetic regulation in active maintenance of β-cell differentiation (Gao et al., 2014) (Friedman and Kaestner, 2006) (Talchai et al., 2012) (Dhawan et al., 2011). Here we add the Polycomb-group protein Eed, and thus the PRC2 complex, to that list. *Eed* knockout, which abrogates both *Ezh1*- and *Ezh2*-dependent catalytic activities as well as H3K27me3 reader capacity, resulted in generalized de-repression of hundreds of bivalent, H3K27me3 marked genes and progressive loss of β-cell identity. Importantly, Eed-knockout triggered transcriptional and pathophysiological changes that recapitulated our evaluations of human and mouse beta-cell dysfunction, namely dysregulation of bivalent and chromatin state A embedded genes, and progressive loss of terminal beta-cell identity. The data highlight the importance of PRC2 for maintaining proper gene expression in both poised and active regions, and indicate that fundamental transcriptional dysregulation is a novel driving feature of beta-cell dysfunction.

Our data highlight a novel role for PRC2 in maintaining functional β-cell identity and importantly, they recapitulate to great extent the chromatin-state associated transcriptional dysregulation observed in mouse and human T2D. In both settings observed dysregulation was characterized by inappropriate and broad activation of progenitor and immature β-cell programs, loss of β-cell function and were consistent with the notion of beta-cell dedifferentiation. The findings that PRC2 loss-of-function triggers a type-2 diabetes-like transcriptional dysregulation and dedifferentiation warrant deeper investigations into relative *Ezh1p*-, *Ezh2*- and non-methyltransferase-dependent contributions of PRC2 in the β-cell compartment. They also beg exploration of more subtle Polycomb-deficient models such as βEedKO-heterozygotes and adult-inducible deficiencies as well as deletions of alternate complex members. Combining these approaches will ultimately define the relative importance of H3K27me2/3 loss, ectopic gene activation and cell turnover towards maintenance of optimal terminal differentiation (Bintu et al., 2016).

Previous reports have indicated glucotoxicity as an important trigger for dedifferentiation (Wang et al., 2014),(Guo et al., 2013). Our findings from insulin-pelleted βEedKO animals, as well as from our mosaic/normoglycemic Pdx-EedKO mice demonstrate that PRC2-associated dedifferentiation does not require hyperglycemia. The data demonstrate that Eed deficiency alone is sufficient to drive terminal dedifferentiation. It will be important now to investigate potential synergistic interactions between PRC2-insufficiency and hyperglycmia. Indeed, we consider a “two-hit” model for dedifferentiation, one in which hyperglycemia and PRC2-insufficiency compound one another to ultimately drive dedifferentiation, as most likely.

The late *down*-regulation and relative loss of active mark enrichment at H3K27ac-broad loci suggest that PRC2 also acts to ensure ‘transcriptional focusing’ at these unique loci. Recent work indicates that spatial confinement is the key parameter controlling frequency of genomic encounters and thus building of super-enhancers and transcription factories (Levine, 2014). Our data are consistent with the idea that H3K27me2/3, which decorates nearly 80% of the genome, enhances 3D compartmentalization of transcriptional regulatory machinery (Boettiger et al., 2016)(Ferrari et al., 2014), and that the highly active subset of chromatin state A embedded genes are particularly dependent on such regulation. Our data add to a select few examples in disparate cell systems where PRC2 actually promotes transcription (Di Croce and Helin, 2013). In addition to its requirement for long-term repression of β-cell progenitor genes upon maturation therefore, PRC2 actively supports terminal differentiation maintenance by also ensuring transcriptional fidelity of chromatin State A embedded lineage-defining regulators.

In summary, the data presented here highlight chromatin state specific transcriptional dysregulation, dedifferentiation and alternate lineage activation as key transcriptional features of beta-cell dysfunction. They demonstrate that Polycomb-mediated gene silencing maintains specificity of the β-cell program by both repression of immature/alternate lineage potential and focussing of transcription at key MβTF’s. The findings indicate that PRC2 functions chronically in β-cells, that PRC2-insufficiency contributes to the pathology of human type-2 diabetes, and therefore, that inhibition of PRC2-opposing factors might serve as potential therapeutic targets for T2D.

### Experimental Procedures

#### ChIP Sequencing

NEXSON ChIP-seq workflow (Arrigoni et al., 2015) is applied to freshly isolated mouse islets. ChIP-seq performed using antibodies against H3K27me3 (Diagenode, #C15410195), H3K9me3 (Diagenode, #C15410193), H3K27ac (Diagenode, #C15410196) and H2AK119ub (Cell signaling, #8240). A list of ChIP-seq data sets used can be found in Table S4.

#### Single-cell and bulk RNA Sequencing

Single-cell RNA-sequencing was performed using CEL-Seq2 method (Hashimshony et al., 2016) with several modifications. For bulk RNA-sequencing, Trizol-purified RNA from mouse islets was poly(A)-enriched and libraries were prepared with a TruSeq Sample Prep v2 kit (Illumina) and sequenced on a HiSeq 2500 (Illumina). Details on bulk and single-cell RNA sequencing can be found Extended Experimental Procedures.

#### Bioinformatic Analyses

For mouse RNA-seq dataset, reads were mapped with TopHat v2.0.13 against mouse genome version GRCm38. For single-cell RNA-seq data, read 2 of each read pair was first 3’ trimmed for adapters, base quality and poly-A tails using cutadapt 1.9.1. Reads were counted with featureCounts (subread-1.5.0-p1) against gene models from Gencode version M9. Differential expression analysis was performed with edgeR (v3.14). For isoform-level analysis we used Isolator (http://www.biorxiv.org/content/early/2016/11/20/088765). Gene set enrichment analysis used GSEA 2.0 with default parameters (Permutation type: gene_set). For human dataset, DESeq2 package v1.8.1 (Love et al., 2014) was used for differential gene expression in diabetic patient samples. ChIP-seq datasets were mapped with Bowtie2 (v2.2.8). We used MACS2 (v2.1.1.20160309) (Zhang et al., 2008) to call peaks over input. Average signal per genesets was calculated using DeepTools v2.4.1 (Ramírez et al., 2016). Analysis of chromatin states used EpiCSeq algorithm (Mammana and Chung, 2015). For all data, statistically significant was adjusted p-value < 0.05. Details of the other analysis can be found in the Extended Experimental Procedures.

#### Human Sample Analysis

Human pancreatic sections were obtained from JDRF nPOD (Network for Pancreatic Organ Donors with Diabetes, see acknowledgement). Sections were stained and quantified using custom Cell Profiler pipeline. Human islet RNA-seq data were obtained from GSE50398 (Fadista et al., 2014) and ChIP-seq data from GSE50386 (Bramswig et al., 2013).

#### Mouse *In Vivo* studies

β-cell specific Eed knockout mice (βEedKO) were generated by crossing *Eed*^*fl*/fl^ mice (Xie et al., 2014) with RIP-Cre transgenic mice. Cre-recombinase-positive wild type and Cre-recombinase-negative *Eed*^*fl/fl and +/fl*^ litter-mates served as controls. Oral glucose (1 g/kg) tolerance test was performed on 8, 16 and 25 weeks old animals after 14 hour overnight fast. Plasma samples were collected for insulin measurements. For HDACi experiment, SAHA was delivered through drinking water as previously described ((Benito et al., 2015)). Mice were treated for 12 weeks from 8 weeks of age before OGTT examination. Animals were kept on a 12 hour light/dark cycle with free access to food and water and housed in accordance with international guidelines. All other mouse models are described in the Extended Experimental Procedures.

#### Histological Studies

Mouse Pancreatic tissue were fixed, paraffinized and sectioned for immunohistochemistry. Images were acquired using Zeiss Apotome or Zeiss LSM 710. Fluorescence intensity, β-cell area and size analysis were performed using semi-automated morphometry using ImageJ or Cell Profiler. Details on staining procedures and analysis pipelines are described in the Extended Experimental Procedures. A list of all antibodies used can be found in Supplementary Table S5.

#### Ectopic Transcription Factor Over-Expression

Min6-B1 Cell lines were electroporated with PMAXGFP (Lonza) or mouse cDNA ORF clones of the 6 different transcription factors (Origene). Cells were harvested after four days post transfection for single-cell RNA-sequencing. Details of the experiment can be found in the Extended Experimental Procedures.

#### Statistical Analysis

Data are expressed as mean ± standard error of the mean (SEM) unless otherwise specified. Statistical analyses were performed as described in the Extended Experimental Procedures. All reported p-values are two-tailed unless stated otherwise, p < 0.05 was considered to indicate statistical significance.

#### Other Methods

See Extended Experimental Procedures.

## Author Contributions

Conceptualization, T.T.-H.L. and J.A.P.; Methodology, T.T.-H.L, S., J.A.P., H.X, S.H.O.; Software, S.H., E.C., D.G; Formal Analysis, Investigation, and Visualization, T.T.-H.L, S.H., E.D., T.B., S., L.L., J.A.P., E.C., C.-H.Y., L.E., K.D., L.A., M.S., U.B., R.T., M.R., S.J.R., F.C.L, B.G.H., A.L.; Writing-Original Draft, T.T.-H.L, J.A.P.; Writing – Review & Editing, T.T.-H.L, J.A.P., A.L., F.C.L., B.G.H.; Supervision, Resources, and Funding Acquisition, J.A.P., T.V., D.G.

## Acknowledgements

S. Kubicek, G. Gradwohl, R. Schneider, T. Jenuwein, A. Öst, K. Gossens, J. Longinotto and S. Welz for discussion, technical and analytical help and advice. This research was performed with the support of the Network for Pancreatic Organ Donors with Diabetes (nPOD), a collaborative type 1 diabetes research project sponsored by the Juvenile Diabetes Research Foundation International (JDRF). Organ Procurement Organizations partnering with nPOD to provide research resources are listed at www.jdrfnpod.org/our-partners.php. This work was supported by funding from the Max-Planck Society, ERC (ERC-StG-281641), BMBF (DEEP), DFG (SFB992 “MedEp”), and EU_FP7 (NoE “Epigenesys”).

---

**Supplementary Figure S1.**
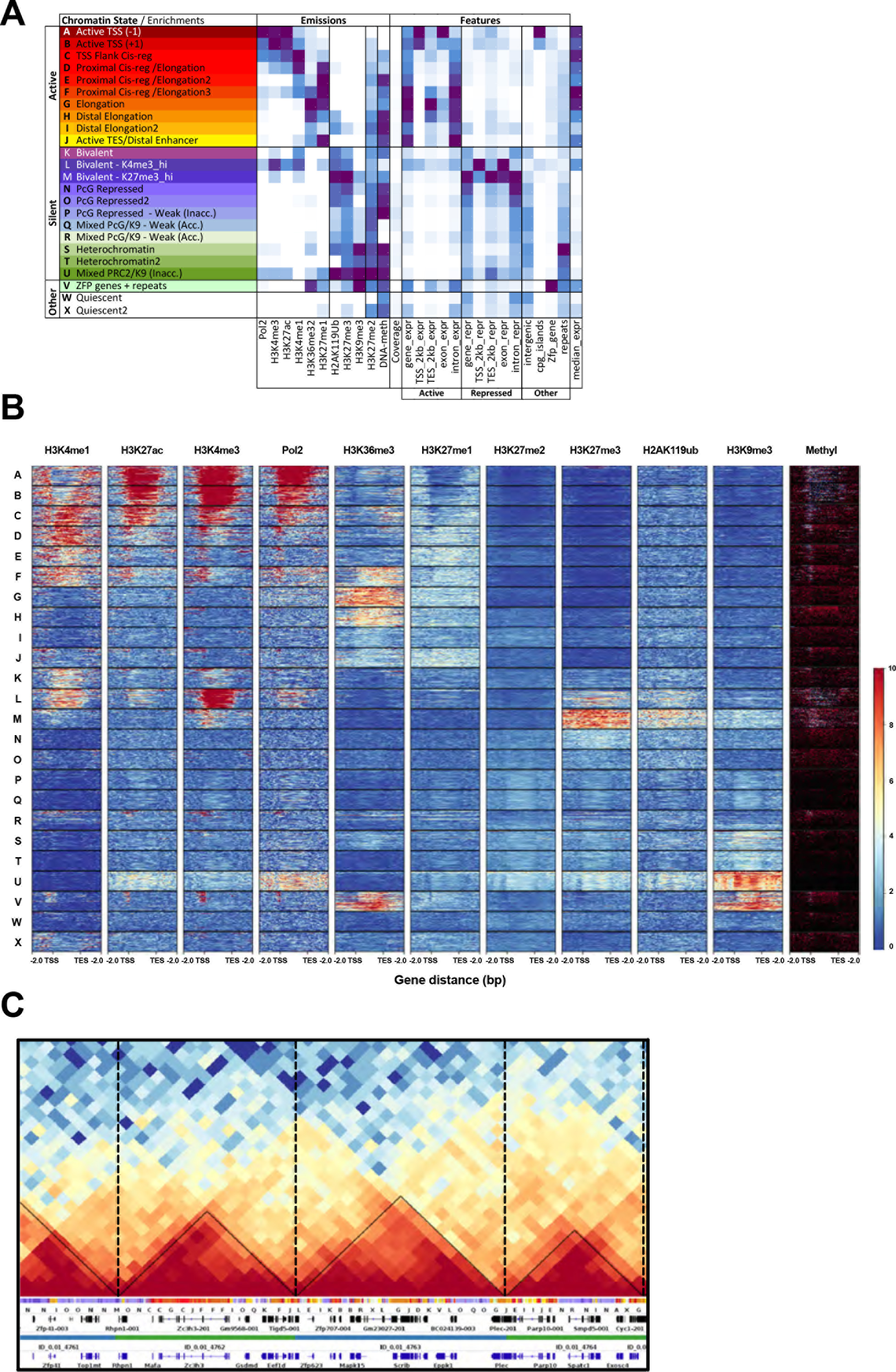
A. Histone mark characteristics, gene expression levels and genomic features of healthy islets. EpiCSeq chromatin state segmentation and manual annotation according to emissions and annotations in D. (Inacc=Inaccessible; + 1 or −1=00B1 nucleosome position around TSS, expr = expressed, repr= repressed, median_expr= Median expression) B. Heatmap profiles of average normalized read coverage from ChIP-seq signals for various histone marks, Pol2 and CpG methylation signal of 100 genes from each chromatin state. C. Example region showing Hi-C TAD boundaries aligning with chromatin state segmentation boundaries. Dash lines indicates TAD boundaries. (Hi-C image taken from http://choroaenome.ie-freibura.mpg.de; dataset mouse CH12; GEO accession GSE63525)

**Supplementary Figure S2.**
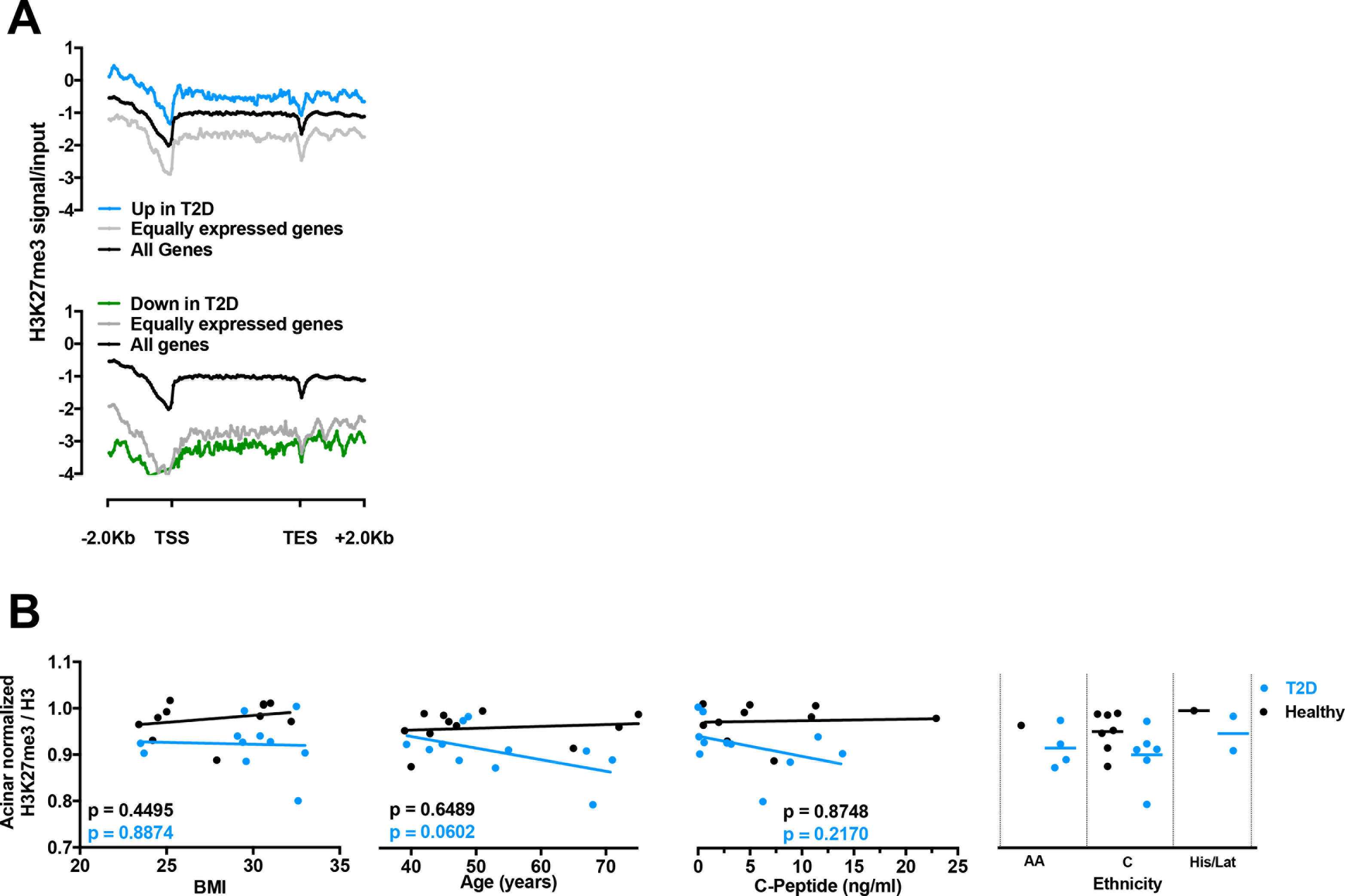
A. Average H3K27me3 signals across genes that are up or down-regulated in T2D, compared to genes that are equally expressed or all genes. TSS: transcription start site, TES: transcription end. B. Reduction of H3K27me3 immunostaining intensity in β-cells of pancreatic sections from T2D donors is independent of BMI, age, C-peptide levels or ethnicity. Solid lines = linear regression analysis of groups. P < 0.05 indicates regression slopes being significantly non-zero.

**Supplementary Figure 3.**
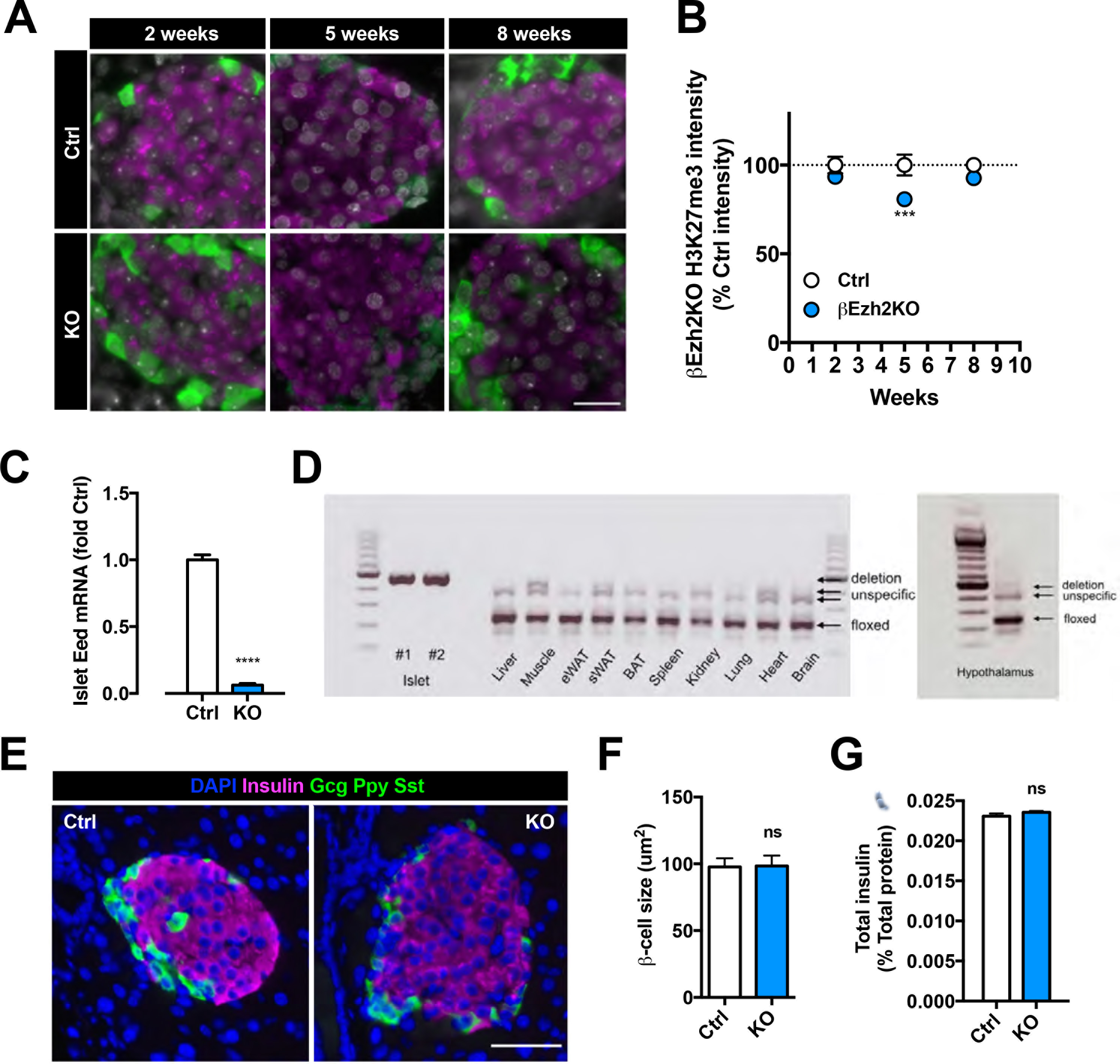
A. Representative images for H3K27me3 staining (grey), insulin (magenta) and glucagon (green) in Ctrl and βEzh2KO at 2, 5 and 8 weeks of age. Yellow arrows indicate β-cell nuclei. B. Relative βEzh2KO β-cell H3K27me3 shows slight reduction relative to Ctrl mice at all time points, with 5 weeks being statistically significant. All data points were normalized to alpha-cell H3K27me3 levels in the same islets prior to comparison. C. Quantitative RT-PCR showing loss of Eed mRNA in βEedKO islets. D. PCR validation of *Eed* allele deletion in islets and other tissues of βEedKO animals. One animal is represented here for all tissues except for islets where two animals are shown E. A faint deletion band of the *Eed* allele can be observed in the hypothalamic extracts of βEedKO animals as expected. F. Immunofluorescence staining of islet hormones shows no dramatic morphological changes between Ctrl and βEedKO islets at 8 weeks of age. G. Quantification of total β-cell mass and insulin area per islet shows no significant difference between Ctrl and βEedKO animals at 8 weeks of age. Data represents mean ± SEM. Scale bar = 25uM. *** p<0.001, **** p<0.0001, ns = p>0.05.

**Supplementary Figure 4.**
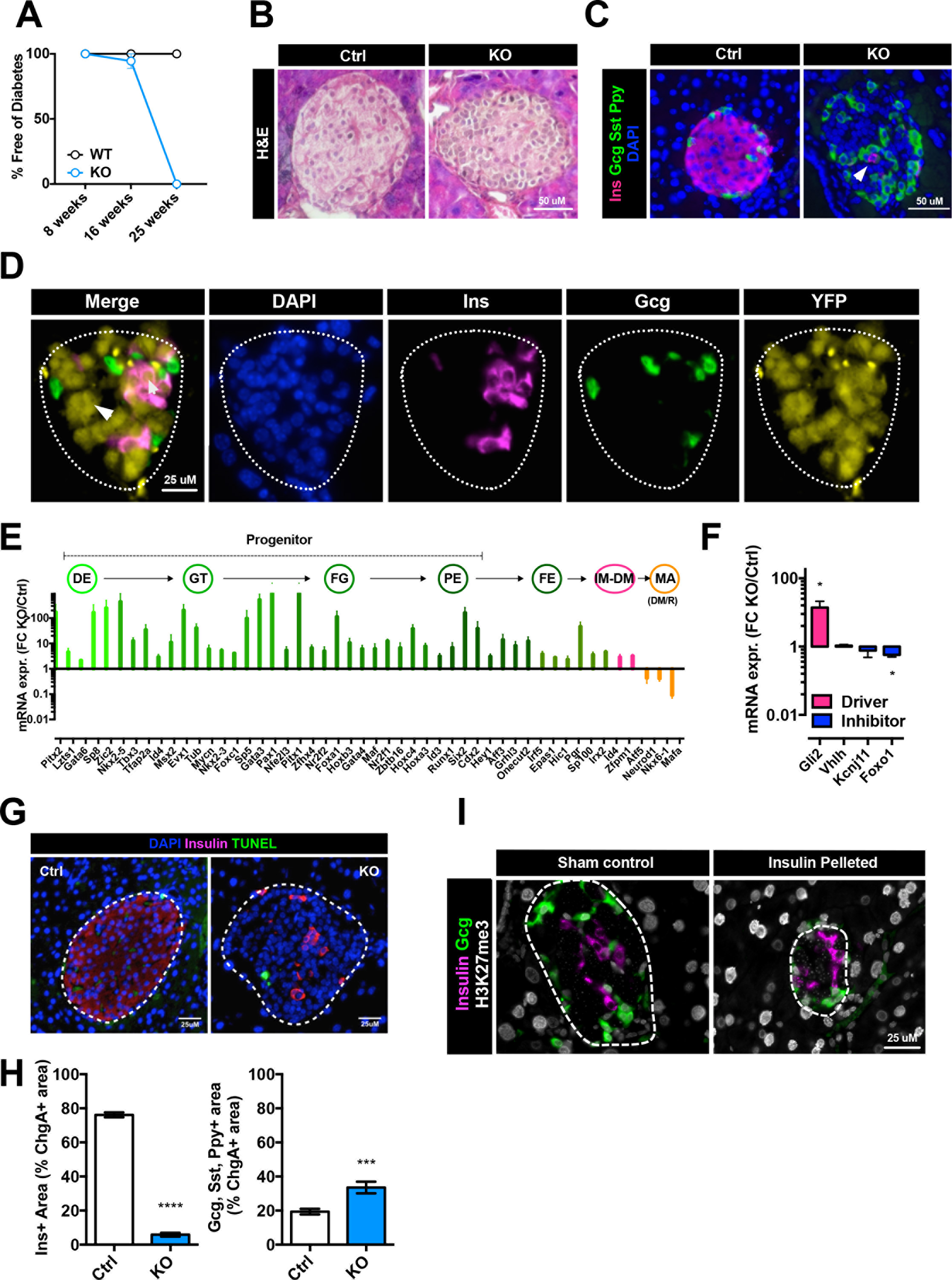
A. Percentage of diabetes-free individuals at 8, 16 and 25 weeks of age. Diabetes was defined as blood glucose >300mg/dL after 2 hours fasting. Note that all animals developed diabetes at 25 weeks of age B. H&E staining of pancreatic sections showing readily detectable islets and reduced inter-nuclei spacing in βEedKO mice C. Immunofluorescence of βEedKO islets shows “empty” cells devoid of staining for major pancreatic hormones and dramatic loss of insulin-producing cells. White arrow indicates residual insulin-positive β-cells. D. Representative image of (βEedK0 islets harbouring a RIP-cre inducible YFP lineage tracer (yellow) in individual channels for clarity. Small white arrow indicates insulin-positive β-cells exhibiting YFP signals, and large white arrow indicates non-insulin expressing cells that also exhibits YFP signal. White dotted line outlines the islet for clarity. E. Mean expression fold change of transcription factors characteristic of developmental precursor stages (DE: definitive endoderm, GT: gut tube, FG: fore gut, PE: pancreatic endoderm, FE: functional endocrine; Xie et al., 2013), immature and mature β-cells (IM-DM/MA-DM; Blum et al., 2012, MA-R: REACTOME: *Regulation of Gene Expression in Beta-Cell*). Arrow represent hypothetical developmental trajectory. All genes listed are core enrichment genes from GSEA analysis and significantly different between βEedKO and Ctrl islets with FDR-q<0.05 F. Gene expression fold change of known factors implicated in β-cell de-differentiation. * FDR-q*0.05 G. Representative images of TUNEL treated Ctrl and βEedKO pancreas showing that dedifferentiated βEedKO β-cells are not apoptotic. White dotted line indicates the border of islets H. A. Morphometric analysis of % β-cell area (Ins+) and α- (Gcg+), δ- (Sst+) and Pp-cell (Ppy+) area within the endocrine compartments of islets at 25 weeks of age. Ins+ cells are less than 5% of the total endocrine area. *** p<0.001, **** p<0.0001 I. Representative images of βEedKO islets from animals treated with insulin pellet or sham controls for 7 weeks from 18 weeks of age.

**Supplementary Figure 5.**
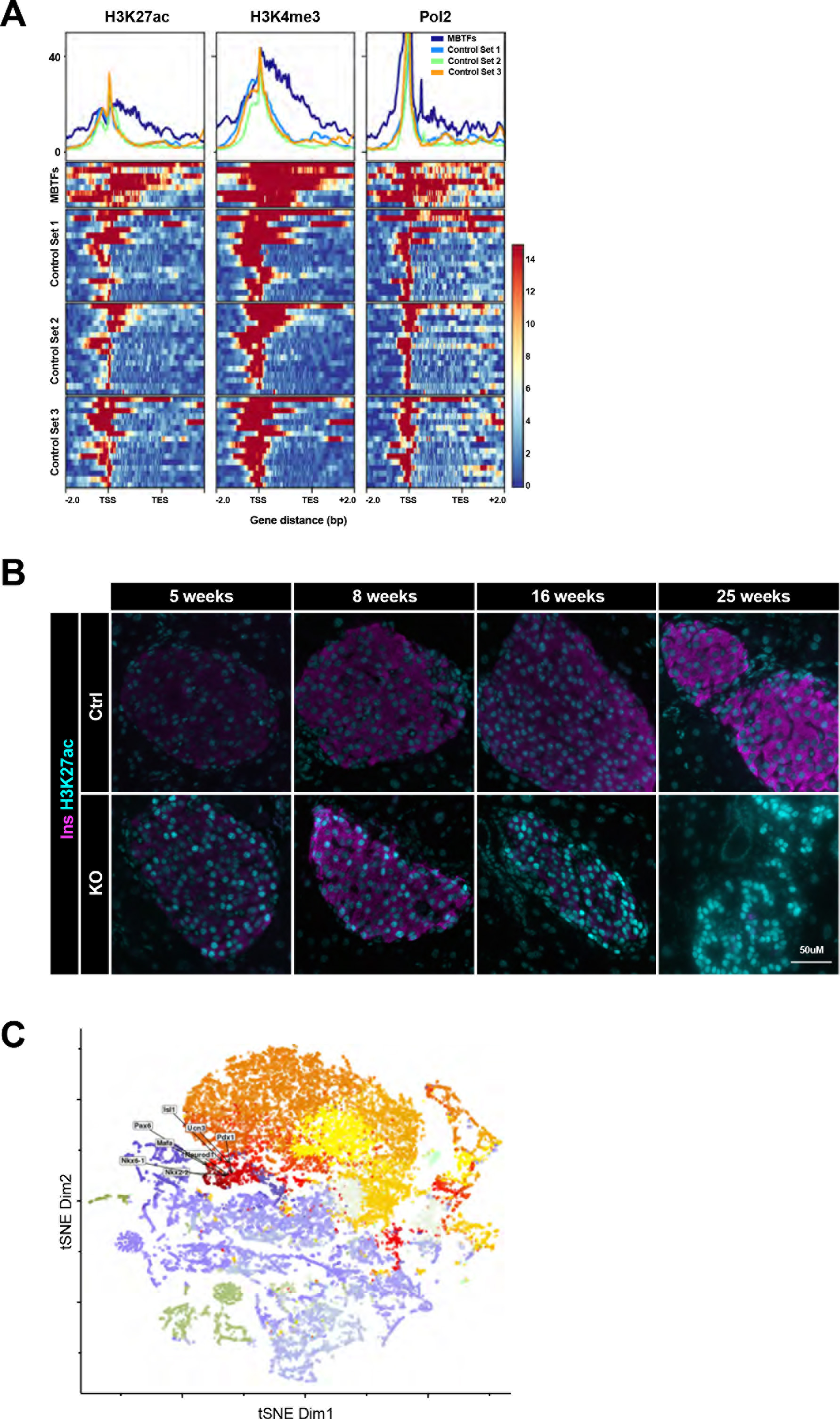
A. Profiles of average normalized read coverage from ChIP-seq signals for H3K27ac, H3K4me3 and Poll I for MβTFs and control gene sets of similar gene expression in mouse β-cells (top). Profiles are presented as heatmaps and sorted according to H3K4me3 signals (bottom). B. Representative images for H3K27ac (cyan) and insulin (magenta) in Ctrl and βEedKO showing progressive gain of H3K27ac in βEedKO β-cells with age. C. tSNE representation all genes in each chromatin states color coded based on Figure 1E. MβTFs highlighted are embedded in the highly-active state A/B (red)

## EXTENDED EXPERIMENTAL PROCEDURES

### Animal Husbandry

All transgenic animals were maintained on a normal chow diet with 15% fat (Ssniff GmbH), fed ad libitum with free access to water (HCl acidified, pH 2.5-3) under controlled humidity and temperature with a 12-hour light and 12-hour dark cycle. Obese mice were fed with high fat diet (60% kcal% fat, Research Diet). All animal studies were performed with the approval of the local authority (Regierungspräsidium Freiburg, Germany) under license number 35.9185.81/G-10/94.

### Generation of βEedKO, βEzh2KO, Pdx1-EedKO and lineage-tracing βEedKO-YFP Mice

Breeding pairs of RIP-cre (Tg(Ins2-cre)23Herr), PDX-1-cre, *Eed*^flox/flox^, Ezh2^flox/flox^ and YFP-reporter (B6.129X1-Gt(ROSA)26Sortm1(EYFP)Cos/J) transgenic mouse line (C57B6/J) were kindly provided by Pedro Herrera, Gerald Gradwohl, Stuart Orkin and Thomas Boehm respectively. To generate βEedKO animals, *Eed*^flox/flox^ animals were crossed with RIP-cre positive *Eed*^+/flox^ animals to obtain mice homozygous for the floxed *Eed* locus. βEzh2KO and Pdx1-EedKO were generated in similar fashion. By crossing animals harboring the YFP-reporter transgene to βEedKO animals, we generated lineage-tracing βEedKO-YFP mice. All mice had been backcrossed for over 10 generations before any phenotyping was initiated.

### Mouse embryo isolation, section preparation and staging

Female mice used for breeding were synchronized for their estrous cycle by exposing them bedding of mature male mice for 48hrs. After setting up the matings, vagina plugs were examined daily and the day of detection was designated E0.5. Pregnant mice were sacrificed on E14.5 and E17.5 using isofluorane and cervical dislocation. Embryos were dissected out of the uterus, washed in 4°C PBS and incised at the cervical region before overnight fixation in HistoFIX solution (Sigma) at 4°C. Embryos were then washed twice for 30 minutes each at 4°C, and dehydrated in 25% EtOH/PBS for 2 hours, 50% EtOH/dH2O for two hours, 70% EtOH/dH2O overnight, 95% EtOH/dH2O for 3 hours and 100% EtOH for 3 hours at 4°C. Embryos were then cleared with xylene for 20 mins each and embedded in paraffin. 5um sagittal sections were collected on SuperFrost glass slides (Fisher). Hematoxylin-Eosin stainings were performed on some sections for confirmatory staging of the embryos. E14.5 embryos were identified for their digitally separated fingers, and absent eyelids. E17.5 embryos were identified for their wrinkled skin and long whiskers upon isolation, performed with reference to <<The Atlas of Mouse Development>> by Kaufman (1992).

### Glucose tolerance test

For the oral glucose tolerance test (OGTT), mice were fasted overnight (16 hrs), whereafter basal blood glucose was measured. Mice were given glucose (1 g/kg) by oral gavage. Blood glucose levels were measured using a OneTouch Vita blood glucose meter at 0, 15, 30, 45 and 60 minutes after glucose. Blood drawn using heparinized capillary tubes was centrifuged at 2000g for 15mins at 4°C. Plasma obtained was used for insulin level measurement by ELISA (Mercodia Ultrasensitive Mouse Insulin Kit).

### Islet isolation, culture and GSIS

Primary islets were obtained by collagenase digestion with reference to (Li et al., 2009). In brief, adult pancreata were perfused through the common bile duct using a 27/30-gauge needle with Collegenase IV (Sigma) solution. Perfused pancreata were digested and purified by centrifugation in Histopaque gradient (Sigma). Isolated islets were hand picked and cultured in complete media (RMPI-1640 containing 11mM glucose, 10% FBS, 0.1% Penincillin/Streptomycin) and maintained at 37°C in 5% CO2 environment.

For GSIS, approximately 30 size-matched and handpicked islets were cultured per well of a 96-well ECM coated plates ( Biological Industries, Israel) and allowed to attached for 48 hours. Islets were then washed twice with Kreb’s buffer with 2.8mM glucose and incubated for 1 hours in Kreb’s buffer with 2.8mM glucose. After incubating islets with another hour of Kreb’s buffer at 2.8mM Glucose (basal), islets were then incubated for 60 mins in Kreb’s buffer supplemented with either 2.8mM Glucose, 11mM glucose or 16.7mM glucose (stimulated). Supernatant fraction for basal and stimulated exposure to glucose were collected, spun for cell removal, and stored −20°C before analysis.

### Total RNA and protein extraction

Purified islets isolated were lysed directly in TRI reagent and total RNA was extracted according to the manufacturer’s instructions. Briefly, cells were lysed in 1ml TRI reagent and lysate was thoroughly mixed by pipetting. Samples in 1.5 ml tubes were incubated for 10 mins at RT and 100ul of 1-bromo-3-chloropropane was added. The mixture was shaken vigorously for 15 seconds and allowed to stand for another 15 minutes at RT. Tubes were centrifuged at 12000g for 15 minutes at 4°C and the aqueous phase was carefully transferred to fresh low-binding tubes (Eppendorf). 500 ul of 2-propanol and 1 ul of GlycoBlue Coprecipitant (Life Technologies) were added to the aqueous phase, mixed and incubated overnight at −20°C before centrifuging them at 12000g for 10 minutes at 4°C. Supernatant was decanted and the RNA pellet washed twice with 1 ml 75% ethanol, air-dried and dissolved in 20 ul of RNase free water (Qiagen). RNA concentrations were quantified on a Qubit 2.0 Fluorometer (Life Technology).

### Histological preparation of the pancreas

Whole pancreata were dissected, weighed, washed in cold PBS and incubated for 24 hours at 4°C in HistoFix (Sigma). For β-cell mass quantification, the pancreata were fixed in homemade cylindrical tube for form standardization of downstream quantification analysis. Each pancreas was then washed twice in PBS for 30 minutes each at room temperature, and dehydrated in 70% EtOH/dH2O overnight, 95% EtOH/dH2O for 3 hours and 100% EtOH for 3 hours at RT before being cleared with xylene for 15 mins and embedded in paraffin. 5 um sections were collected on SuperFrost glass slides (Fisher).

### Immunofluorescence, image acquisition and analysis

Paraffinized sections (mouse and human) were prepared for immunofluorescence staining by heating the slides for 15 mins at 55°C in an oven, deparaffinized (2 × 100% xylene 5 minutes each, 2 × 100% ethanol 5 minutes each, 2 × 95% ethanol 5 minutes each, 70% ethanol for 5 mins) and rinsed in dH2O for 5 minutes. Antigen retrieval was performed by heating the slides at 95°C for 20 minutes in HistoVT pH 7.0 (Nacalai USA) for all antibodies used. Specimens were blocked in 5% goat serum PBS-T for 15 minutes at RT before incubating with primary antibody diluted in 1% goat serum PBS-T overnight at 4°C. For primary antibodies produced in goat, donkey serum was used as the blocking agent. Each slide was rinsed three times in PBS, for 5 minutes each. Specimens were incubated in fluorochrome-conjugated secondary antibody diluted in 1% goat serum PBS-T for 1 hr at RT in the dark. After rising as above, VectorShield with DAPI and coverslip were mounted and slides were allowed to cure overnight at 4°C in the dark before image acquisition. For apoptosis assessment, we used DeadEnd Fluorometric TUNEL (Promega) and performed the staining as recommended by manufacturer. A list of all antibodies can be found in Supplementary Table 6.

Images were acquired using either ApoTome.2 (Zeiss) without structured illumination or LSM780 (Zeiss) and analyzed using ImageJ software. For intra-islet quantification, 15-30 islet were chosen randomly from at least 2 sections spaced ~200 um apart for each animal. 4-5 animals were used per genotype. For mouse samples, H3K27me3 positivity and intensity were assessed using Image J measurement functions in a blinded fashion. For human samples, analysis of H3K27me3, H3, Insulin and DAPI intensities were automated using customized Cell Profiler pipeline (available upon request).

### β-cell, α-cell mass quantification

For all morphometric analysis of islets, 4-5 animals of each genotype were analyzed. Each section was stained for insulin (β-cells), glucagon (and pancreatic polypeptide and somatostatin) and/or Chromogranin A (total islet area) and DAPI (cell count and total pancreas area estimation). For β-cell mass analysis, 4 sections ~200 um apart were covered systematically by accumulating images from non-overlapping fields with ApoTome.2 (Zeiss) using a 10X objective for whole pancreas section. For other endocrine cell area analysis, 20-50 islet profiles were chosen randomly from each sections and captured using a 40X objective. All morphometric analyses were performed using ImageJ software. Briefly, individual channels were converted to 8-bit grayscale and measurement scale was converted from pixels to μm. A identifcal threshold was applied to all images from the same channel to exclude background signals and further converted to binary format before automated analysis of immunoreactive area. β-cell area was expressed as a fraction of total surveyed pancreatic area and β-cell mass was estimated by multiplying the fraction to pancreas weight. Other endocrine cell area was expressed as a fraction of total islet area surveyed by total islet area from Chromogranin A immunoreactivity. β-cell size was estimated by dividing the total β-cell area by the total number of β-cell nuclei that it contained. For all analysis, the region of interest was traced carefully to exclude other tissues such as ducts, lymph nodes and mesenteric tissues.

### Bulk RNA-seq and bioinformatic analysis

mRNA from whole islets was used to generate libraries using Illumina TruSeq RNA Sample Prevp v2 (RS-122-2001). The manufacturer’s recommendations were followed and the libraries were sequenced on an Illumina HiSeq 2500 sequencer. All sequence data were performed in biological triplicates, each containing islets from 1-2 animals, at 2 × 50 bp length with high quality metrics (>20 Phred score) and nucleotide distribution. For 25 weeks βEedKO samples, two biological replicates were used, each containing islets from at least 4 animals.

Reads for mouse bulk RNA-seq datasets were mapped with TopHat v2.0.13 against mouse genome version GRCm38. The total number of sequenced reads ranged from 10-50 million pairs of which at least 68% of the reads were mapped. Reads were counted with featureCounts (subread-1.5.0-p1) against gene models from Gencode version M9. Differential expression analysis was performed with edgeR (v3.14). Alignment statistics (data not shown) indicated data were of high quality and sequencing depth was sufficient to test for differential expression between conditions. Isoform-level analysis was performed with Isolator (http://www.biorxiv.org/content/early/2016/11/20/088765). For human dataset, DESeq2 package v1.8.1 (Love et al., 2014) was used for differential gene expression in diabetic patient samples. The count matrix was used as given (GEO: GSE50244). Differential genes were called with an FDR threshold of 0.05. The mapping from mouse to human orthologues was done with Biomart (http://www.ensembl.org/biomart). Correlations between mouse and human expression fold changes were calculated in R using Pearson correlation.

### Transcription factor over expression and Single Cell RNA-seq

Min6-B1 cell line were electroporated using Nucleofector^®^ II device (Lonza, Kit T, Program 0-017) with pMAXGFP (Lonza) or mouse cDNA ORF clones of the 6 different transcription factors (Origene). 4 days after electroporation, cells were harvested, trypsinized and single cells were sorted in 384-well plates containing 240nL of primer mix and 1.2uL of PCR encapsulation barrier, Vapor-Lock (Qiagen GmbH, Germany).

Single cell RNA sequencing was performed using CEL-Seq2 method (Hashimshony et al., 2016) with several modifications. Importantly, a fivefold volume reduction was achieved using a nanoliter-scale pipetting robot, Mosquito HTS (TTP Labtech). Sorted plates were centrifuged at 2200g for 10min at 4°C, snap-froze in liquid nitrogen and stored at −80°C until processed. 160nL of reverse transcription reaction mix and 2.2μl of second strand reaction mix was used to convert RNA into cDNA. cDNA from 96-cells was pooled together before clean up and in vitro transcription, generating 4 libraries from one 384-well plate. 0.8μl of AMPure/RNAClean XP beads (Beckman Coulter GmbH, Germany) per 1μl of sample were used during all the purification steps including library cleanup.12 libraries (1152 single cells) were sequenced in a single lane (pair-end multiplexing run, 100bp read length) of Illumina HiSeq 2500 sequencing system generating 200 million sequence fragments.

### Single Cell RNA-seq analysis

Read 2 of the each read pair was first 3’ trimmed for adapters, base quality and poly-A tails using cutadapt v1.9.1. Remaining reads were mapped with STAR v2.5.3.a against mouse genome version GRCm38 with gene models from Gencode version M9. Gene summarization was done using featureCounts (subread-1.5.0-p1) (Liao et al., 2014) collapsing exons to genes. Genes with a biotype related to pseudogenes as well as multimapping reads were discarded. Cell demultiplexing was done using the cell-barcode and the unique molecular identifier (UMI) present in the first 12 nt of Read 2 of the read pair.

Data analysis was performed using RaceID2 and StemID algorithm (Grün et al., 2016). Downsampling to 5,000 transcripts was used for data normalization. K-medoids clustering was performed using 1-Spearman’s correlation as a distance metric. The minimum suitable cluster number (=4) characterizing the dataset was determined by computing Jaccard’s similarity for each cluster by bootstrapping for k-medoids clustering with different cluster numbers. The minimum number yielding a Jaccard’s similarity >0.6 for all clusters was selected. The t-distributed stochastic neighbor embedding (t-SNE) algorithm was used for dimensional reduction and cell cluster visualization (Maaten and Hinton, 2008). Since outlier identification is beyond the scope of the paper, RaceID2 was executed with the probability threshold value for outlier identification set to zero. The StemID algorithm was used to infer a dedifferentiation trajectory. A p-value threshold of 0.05 was chosen to assign significance to the links. StemID identified a dedifferentiation trajectory along the clusters 1-2-3-4.

To identify modules of co-expressed genes along the dedifferentiation trajectory, all cells assigned to these links were assembled in a pseudo-temporal order based on their projection coordinate. All genes that were not present with at least three transcripts in at least a single cell were discarded from the subsequent analysis. The pseudo-temporal gene expression profiles of all genes were subsequently z-score transformed and topologically ordered by computing a one-dimensional self-organizing map (SOM) with 1,000 nodes. We then defined modules of co-expressed genes by grouping neighboring nodes of the SOM if average gene expression profiles at these nodes exhibit a Pearson’s correlation coefficient larger than 0.9. Only modules with more than 5 assigned profiles were retained for visualization of coexpressed genes.

### Gene-Set Enrichment Analysis

For gene-set enrichment analysis, javaGSEA desktop application from Broad institute was used. Cufflink transcriptome profile for each sample was used without any data pre-filtering. Pair-wise gene-set enrichment analysis was performed using Gene Ontology (GO) gene set collection derived from gene2go annotation data from NCBI with the latest update (April_2015), c2cp and manually curated gene sets from (Xie et al., 2013) and (Blum et al., 2012). Number of permutations were set to default 1000 and permutation type set to “gene_set” and chip platform set to “Gene_Symbol.chip”. Enrichment statistics, metric for ranking genes, gene list sort and ordering mode were kept at default values.

### ChIP-sequencing and bioinformatics analysis

NEXSON ChIP-seq workflow (Arrigoni et al., 2015) was applied to freshly isolated mouse islets. All mouse Chip-seq data (our data as well as external data) was mapped with Bowtie2 (v2.2.8) and filtered for PCR duplicates. Bigwig tracks for visualization in IGV were created with DeepTools v2.4.1 and normalized to sequencing depth (Ramírez et al., 2016). For Figure 7, H3K27ac and H3K4me3 peaks were called in “broad” mode in MACS2 (v2.1.1.20160309) (Zhang et al., 2008). Histone breadth is defined as peak length from the start to the end of each called peak regions. A list of ChIP-seq data sets used can be found in Supplementary Table 7.

### Chromatin Segmentation

We used EpicSeg for chromatin segmentation (Mammana and Chung, 2015). We combined our own ChIP-seq data for H3K4me3, H3K27ac, H3K27me3, H3K27me2, H3K27me1, H3K36me3, H2AK119Ub, H3K9me3 and Pol2 together with external available data for H3K4me1, H3K27me3 and H3K9me3 marks (Hoffman et al., 2010) (Tennant et al., 2013). EpicSeg uniquely assigns reads or fragments to genomic bins (size 200bp) and hence is able to combine single and paired end data. Replicates are averaged. In addition we incorporated methylation data from young islets (Avrahami et al., 2015) by adding an additional column to the internal matrix of EpicSeg. For this purpose we calculated the median methylation in a 600bp window and a shift size of 200bp. The binned methylation percentages were then inversely scaled. By this transformation, most of the genomic bins for methylation have low or 0 values (ie. high methylation) whereas regions with a lower methylation have higher values. Chromatin states were assigned to genes according to 1) the maximum single state coverage over the genebody (genebody state) and 2) the chromatin state at the TSS (TSS state). Assignments were only done to the “basic” subset from Gencode M9 and to genes with biotype “protein_coding”,”lincRNA” or “antisense”. Of note, original methylation data (GEO: GSE68618) from young islets (Avrahami et al., 2015) was remapped to GRCm38 with bwa-meth and methylation levels were obtained with MethylDackel/PileOMeth (https://github.com/dpryan79/MethylDackel).

### Statistical Analysis

All data unless otherwise stated are shown as mean value 00B1 standard error of the mean (SEM) and tested statistically using two-tailed Student’s t-test or ANOVA. All figures and statistically analyses were generated using GraphPad Prism 5. p<0.05 was considered to indicate statistical significance.

